# Polyamine metabolism regulates the T cell epigenome through hypusination

**DOI:** 10.1101/2020.01.24.918094

**Authors:** Daniel J. Puleston, Francesc Baixauli, David E. Sanin, Matteo Villa, Agnieszka Kabat, Marcin M. Kamiński, Hauke Weiss, Katarzyna Grzes, Lea Flachsmann, Cameron S. Field, Michal Stanckzak, Lena Schimmelpfennig, Fabian Hassler, Chao Wang, Nir Yosef, Vijay K. Kuchroo, Yaarub Musa, Gerhard Mittler, Joerg M. Buescher, Stefan Balabanov, Edward J. Pearce, Douglas R. Green, Erika L. Pearce

## Abstract

We report here a central role for polyamines in T cell differentiation and function. Deficiency in ornithine decarboxylase (ODC), a critical enzyme for polyamine synthesis, resulted in a profound failure of CD4^+^ T cells to adopt correct subset specification, underscored by ectopic expression of multiple cytokines and lineage-defining transcription factors across T_H_1, T_H_2, T_H_17, and T_reg_ polarizing conditions, and enhanced colitogenic potential. T cells deficient in deoxyhypusine synthase (DHPS) or deoxyhypusine hydroxylase (DOHH), which sequentially utilize polyamines to generate hypusine, phenocopied *Odc*-deficient T cells, and mice in which T cells lacked *Dhps* or *Dohh* developed colitis. Polyamine-hypusine pathway enzyme deficiency caused widespread chromatin and transcriptional dysregulation accompanied by alterations in histone methylation, histone acetylation, and TCA cycle metabolites. Epigenetic modulation by 2-hydroxyglutarate, or histone acetyltransferase inhibition, restored CD4^+^ T cell subset specification. Thus, polyamine synthesis via hypusine is critical for maintaining the epigenome to focus T_H_ cell subset fidelity.

## INTRODUCTION

Upon activation, T cells proliferate to form a population of effector cells that mediate immunity. For CD4^+^ helper T (T_H_) cells, this clonal expansion is linked to their differentiation into distinct subsets of cells with specialized functions. CD4 T_H_ cell differentiation is a key feature of the adaptive immune response and is critical for controlling pathogens and maintaining tissue homeostasis. The functional specialization of T_H_ cells is conferred by the expression of T cell subset-specific transcription factors that coordinate genetic programs to direct production of soluble proteins and surface molecules that allow interactions with other cells (Murphy and Stockinger, 2010).

Three major subsets of effector T_H_ cells include T_H_1, T_H_2, and T_H_17 cells, which are defined by the expression of canonical cytokines such as IFN-γ, IL-5 and/or IL-13, and IL-17, as well as the lineage specific transcription factors T-bet, GATA3, and RORγt, respectively (Murphy and Stockinger, 2010). A fourth subset, regulatory T (T_reg_) cells, is critical for modulating immunity by dampening effector T cell activation and proliferation, and expresses the lineage specifying transcription factor Foxp3 (Fontenot et al., 2003). These four subsets are induced by distinct conditions both *in vivo* and *in vitro*. For the latter, well-established methods exist to polarize CD4^+^ T cells into subsets with high fidelity, where the majority of cells in culture will largely adopt lineage specific traits of their respective identity. These cultures mimic what occurs *in vivo*, where appropriate T cell activation and faithful subset commitment is critical for productive and healthy immunity.

We are interested in the role of metabolic reprograming in T cell activation and differentiation (Buck et al., 2015). Numerous older reports have shown a direct link between polyamines, low molecular weight aliphatic polycations, and T cell proliferation (Bowlin et al., 1987; Kay and Pegg, 1973; Schall et al., 1991; Scott et al., 1985). In mammalian cells, polyamines consist of putrescine, spermidine, and spermine. More recently it was shown that increased levels of polyamines can be detected in T cells after activation, and that pharmacological inhibition of ODC, the first and rate-limiting enzyme in the polyamine synthesis pathway, which converts ornithine to putrescine, inhibits activation-induced proliferation (Wang et al., 2011). While these data highlight the importance of polyamines for T cell clonal expansion, the mechanistic basis for these effects has yet to be elucidated.

In eukaryotes, spermidine serves as a substrate for the hypusination of a conserved lysine residue in eukaryotic initiation factor 5A (eIF5A) (Park et al., 1981) (Park et al., 1981), through the action of DHPS and DOHH. EIF5A is a factor that controls translation elongation and termination (Dever et al., 2018; Saini et al., 2009), and its post-translational hypusine modification by DHPS and DOHH is critical for its function (Park et al., 2010; Park and Wolff, 2018). The only known function of DHPS and DOHH is, via a two-step process, to convert this one lysine residue in eIF5A to the amino acid hypusine (Park et al., 2010). Whole body knockouts of DHPS, DOHH, or eIF5A are embryonic lethal (Pällmann et al., 2015; Sievert et al., 2014). However, polyamines are recognized to have pleiotropic functions (Igarashi and Kashiwagi, 2000; Puleston et al., 2017), and the biological contexts in which polyamine synthesis is coupled to hypusine are not well understood.

In this study, we aimed to understand the role of polyamine metabolism in CD4^+^ T cell differentiation and function. Our data indicate that the loss of function of ODC, DHPS, or DOHH leads to profound changes in the ability of CD4^+^ T cells to faithfully differentiate into functionally distinct subsets. This reflects major changes in the epigenome and transcriptome of activated T cells linked to linked to alterations in core TCA cycle metabolism and chromatin modifier expression. Our data place the polyamine-hypusine axis in a position of central importance in CD4^+^ T cell differentiation.

## RESULTS

### Polyamine biosynthesis via Odc regulates CD4^+^ T cell subset fidelity

Synthesis of the polyamines putrescine, spermidine, and spermine requires ODC (**Fig.1A**), and its expression (**Fig. 1B**), and polyamines (**Fig. 1C**), are increased after CD4^+^ T cell activation. To test the role of polyamine synthesis in CD4^+^ T cells, we bred mice with loxP flanked exons 9-11 of *Odc* with mice expressing *CD4*^*cre*^, to generate mice with *Odc* specifically deleted in T cells (*Odc*-ΔT mice). Control mice were absent for cre recombinase. Efficient deletion of ODC in T cells was confirmed by immunoblot (**Fig. 1D**). As we predicted, *Odc*^*-/-*^ T cells exhibited reduced polyamines after activation *in vitro* (**Fig. 1E**). To explore the significance of polyamine synthesis in activated T cells, we isolated naïve CD4^+^ T cells from *Odc*-ΔT mice and polarized them into T_H_1, T_H_2, T_H_17, and regulatory T (T_reg_) cell subsets (Zhu et al., 2010). After 4 days we restimulated cells with PMA/ionomycin and measured intracellular cytokines. Irrespective of polarization condition, *Odc*^*-/-*^ T cells in each subset displayed significantly elevated IFN-γ production and an increased frequency of cells that produced both IFN-γ and IL-17A (**Fig. 1F**) and IL-17F (**not shown**). Further, there was an increased frequency of IL-5 and IL-13 double producing cells, in both the T_H_1 and T_H_2 subsets (**Figs. 1G** and **S1A**), as well as an increase in IFN-γ and IL-13 double producing cells across all T_H_ subsets (**Figs. 1G** and **S1B**). Of note, while the frequency of IL-17 producing cells increased in T_H_1, T_H_2, and T_reg_ cultures, the frequency of these cells decreased under T_H_17 conditions (**Fig. 1F**). Together these data collectively showed that in the absence of *Odc*, T_H_ cells express dysregulated, and sometimes noncanonical, cytokines, even when activated in polarizing conditions that normally direct T_H_ cell subset specification with high fidelity.

**Figure 1.**
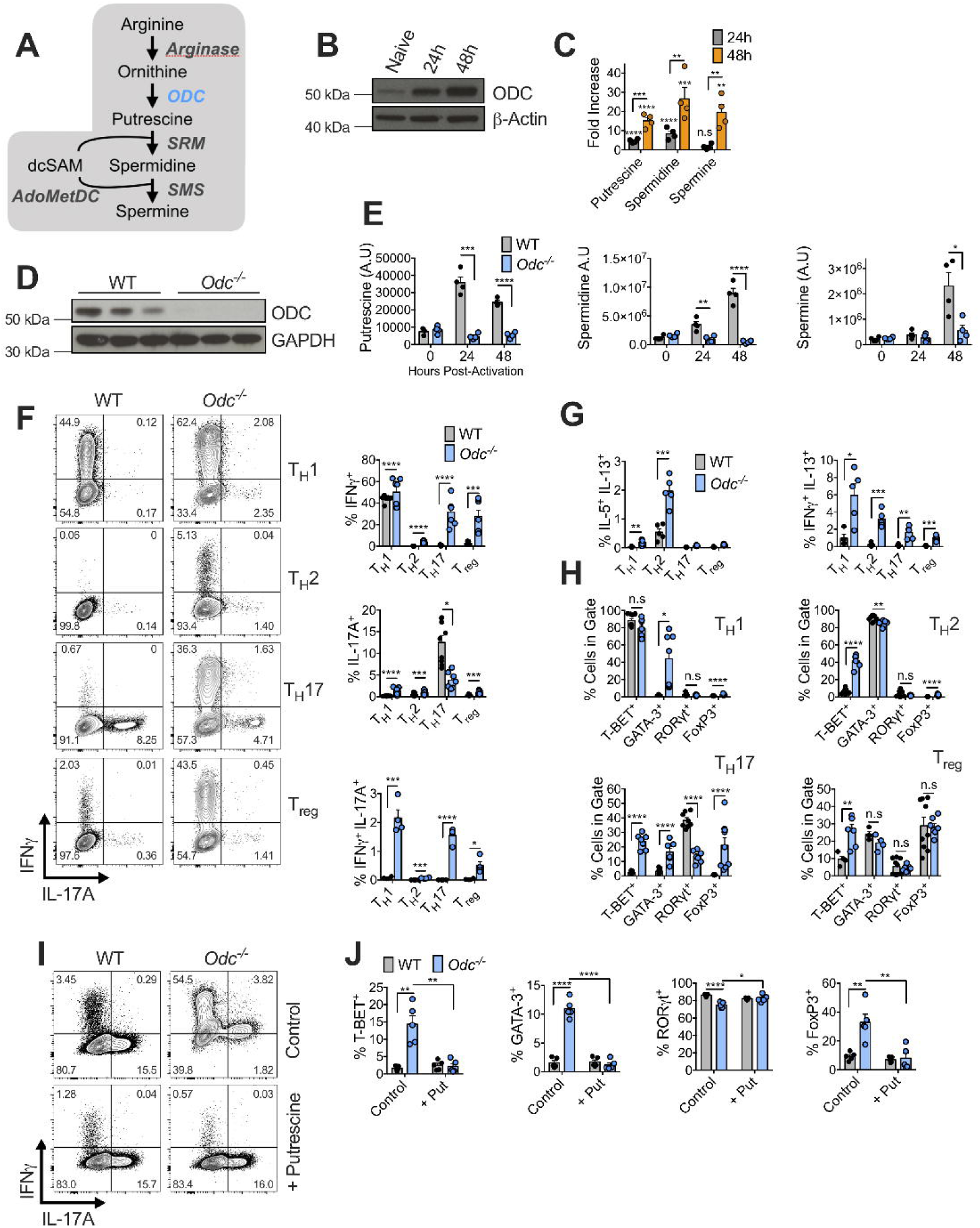
Polyamine biosynthesis via *Odc* regulates CD4^+^ T cell subset fidelity. **(A)** Schematic of the polyamine biosynthesis pathway. (**B**) Immunoblot of CD4^+^ T cells isolated from spleens of WT mice and activated for indicated length of time with anti-CD3/CD28 or rested overnight in 10 ng/mL IL-7, representative of 3 biological replicates. (**C**) Mass spectrometry analysis of polyamine metabolite levels in CD4^+^ T cells following activation with anti-CD3/CD28. Fold increase is relative to unactivated CD4^+^ T cells rested overnight in IL-7. (**D**) Immunoblot of CD4^+^ T cells isolated from spleens and lymph nodes of WT and *Odc*-ΔT mice and activated for 48 hours with anti-CD3/CD28. (**E**) Mass spectrometry analysis of polyamine metabolite levels in WT and *Odc*^-/-^ CD4^+^ T cells following activation with anti-CD3/CD28. (**F** and **G**) WT and *Odc*^-/-^ naïve CD4^+^ T cells were activated with anti-CD3/CD28 for 4 days in indicated T_H_ cell polarizing conditions. Analysis of intracellular cytokine expression 5 hours after PMA/ionomycin re-stimulation by flow cytometry is shown with representative contour plots. (**H**) WT and *Odc*^-/-^ CD4^+^ T cells activated and polarized as in (F) and analyzed for transcription factor expression on day 4 by flow cytometry. (**I** and **J**) WT and *Odc*^-/-^ naïve CD4^+^ T cells activated under T_H_17 cell polarizing conditions ± 250 μM putrescine for 4 days. Intracellular cytokine was assessed 5 hours post re-stimulation with PMA/ionomycin and representative flow cytometry plots are shown (**I**). All data are mean ± SEM (p*<0.05, p**<0.005, p***<0.0005, p****<0.00005). (C, E, I, J) representative of 2 experiments, (F-H) representative of 5 experiments.

In addition to measuring cytokines in *Odc*^*-/-*^ T_H_ cells we also measured lineage-specific transcription factor protein expression. Expression of T-bet, the T_H_1 lineage specifying transcription factor (Szabo et al., 2000), was aberrantly increased in T_H_2, T_H_17, and T_reg_ cells, while GATA3, the T_H_2 lineage specifying transcription factor, was increased in T_H_1 and T_H_17 cells (**Figs. 1H** and **S2A**). While expression of T-bet, GATA3, and Foxp3, the T_reg_ cell specifying transcription factor, were increased in T_H_17 cells, expression of RORγt, the T_H_17 specifying transcription factor, was decreased within the T_H_17 cell culture (**Figs. 1H** and **S2A**). To test whether the phenotype of *Odc*^*-/-*^ T cells resulted from an inability of these cells to synthesize putrescine, we treated control and *Odc*^*-/-*^ T_H_17 cells with putrescine and measured cytokine and transcription factor expression. Importantly, providing exogenous putrescine to the cells restored expression of cytokines and transcription factors to the level of control WT T_H_17 cells (**Figs. 1I, 1J, S2B**, and **S2C**). Together these results showed that *Odc* is critical for enforcing the fundamental programming that underlies T_H_ cell subset lineage specification.

#### Purified naïve Odc^-/-^ CD4^+^ T cells are highly colitogenic in a T cell transfer model of colitis

The *Odc*-ΔT mice appeared grossly normal in terms of overall health and peripheral T cell numbers in the steady state (not shown), but when activated *in vitro, Odc*^*-/-*^ T cells displayed profound cytokine and transcription factor dysregulation (**Figs. 1** and **S1**). We wondered whether activating these cells *in vivo* would reveal an enhanced inflammatory phenotype marked by cytokine and transcription factor dysregulation. To explore this we used a mouse adoptive T cell transfer model of inflammatory bowel disease, where adoptively transferring naïve CD4^+^ T cells into RAG1-deficient recipient mice, which lack their own lymphocytes, drives colitis after approximately 1-2 months (Powrie et al., 1993). We transferred 4×10^5^ naïve (CD45RB^hi^, CD25-, CD44^lo^, CD62^hi^) CD4^+^ T cells isolated from *Odc* WT and *Odc*-ΔT mice into age and sex matched *Rag1*^*-/-*^ recipient mice. After 3 weeks, the *Rag1*^*-/-*^ mice that had received *Odc*^*-/-*^ T cells began to lose weight, which steadily declined until 36 days post T cell transfer when the experiment had to be terminated due to excessive weight loss and the appearance of severe diarrhea (**Figs. 2A** and **2B**). Although colon length was comparable between RAG1-deficient mice that received either WT (WT → *Rag1*^*-/-*^) or *Odc*^*-/-*^ T cells (*Odc*^*-/-*^ → *Rag1*^*-/-*^) (**Fig. 2C**), only the mice containing *Odc*^*-/-*^ T cells showed macroscopic signs of inflammation, characterized by thickening of the colonic walls, a hallmark of colon inflammation (**Fig. 2D**). When we assessed the T cell cytokine profile of *Rag1*^*-/-*^ recipient mice, there was an increased number of T cells expressing IFN-γ in both the colon (**Fig. 2E**) and mesenteric lymph nodes (MLN) (**Fig. S3A**) of *Odc*^*-/-*^ → *Rag1*^*-/-*^ mice relative to WT → *Rag1*^*-/-*^ controls, while the number of IL-17A-, IL-17F- and IL-17A/IL-17F-expressing cells was reduced in the colon, and frequency in the MLN, in *Odc*^*-/-*^ → *Rag1*^*-/-*^ mice (**Figs. 2E** and **S3B**). When we analyzed the expression of transcription factors key to T_H_ lineage commitment, there was a higher frequency of T cells expressing T-bet in the colon of *Odc*^*-/-*^ → *Rag1*^*-/-*^ mice compared to WT controls, while the frequency of T cells expressing RORγt was reduced in *Odc*^*-/-*^ → *Rag1*^*-/-*^ mice (**Fig. 2F**). The frequency and number of T cells expressing Foxp3 and CD25, markers of T_reg_ cells, in the colon and MLN was comparable between WT → *Rag1*^*-/-*^ and *Odc*^*-/-*^ → *Rag1*^*-/-*^ mice (**Figs. S3C** and **S3D**). These data confirmed a critical role for ODC in ensuring correct T_H_ lineage fidelity both *in vitro* and *in vivo*.

**Figure 2.**
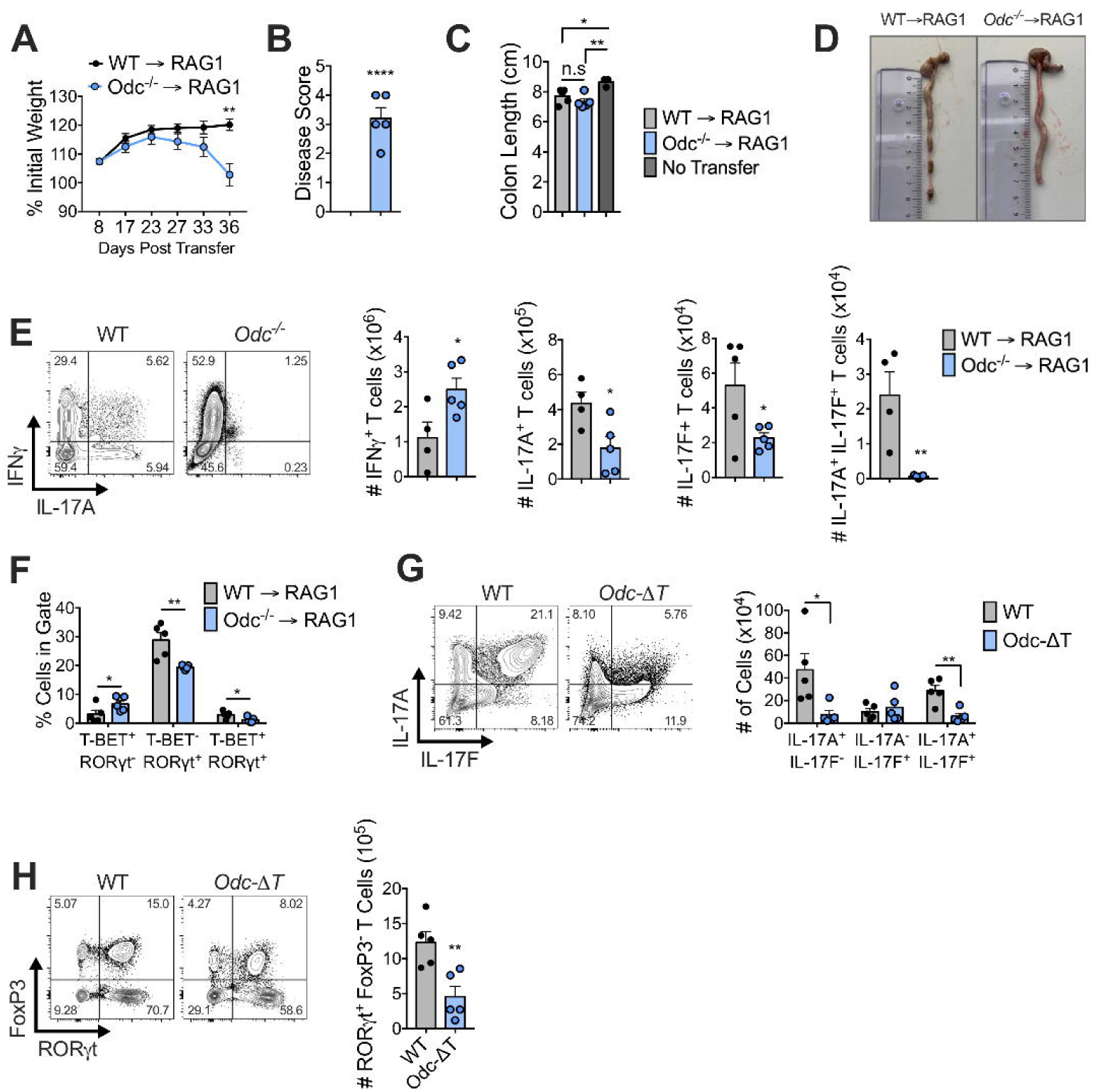
Naïve *Odc*^*-/-*^ CD4^+^ T cells are highly inflammatory in a T cell transfer model of colitis. (**A**) 4×10^5^ WT or *Odc*^*-/-*^ naïve (CD45RB^hi^ CD25^-^ CD44^lo^ CD62L^hi^) CD4^+^ T cells were adoptively transferred into *Rag1*^*-/-*^ recipient mice and weight loss tracked over 37 days. (**B**) Disease score based on weight loss, stool consistency, and presence of blood in stool (see methods) was assessed in *Rag1*^*-/-*^ recipients following adoptive transfer of either WT or *Odc*^*-/-*^ naïve CD4^+^ T cells. Colon length (**C**) and representative images of colons (**D**) from *Rag1*^*-/-*^ recipient mice following adoptive transfer of either WT or *Odc*^*-/-*^ naïve CD4^+^ T cells. (**E**) Number of CD4^+^ T cells expressing indicated cytokine from colon following 4 hours *ex vivo* re-stimulation with PMA/ionomycin, representative contour plots are shown. (**F**) Frequency of colonic CD4^+^ T cells from *Rag1*^*-/-*^ recipient mice expressing indicated transcription factor(s). (**G**) WT and *Odc*-ΔT mice were treated with 50 μg of anti-CD3 antibody by intraperitoneal injection on day 0, day 2 and day 4 and sacrificed 4 hours after the last injection. The number of CD4^+^ T cells expressing indicated cytokine was assessed from the small intestine by flow cytometry following 4 hours *ex vivo* re-stimulation with PMA/ionomycin, representative contour plots are shown. (**H**) CD4^+^ T cells from the small intestine of WT and *Odc*-ΔT mice, treated as in (**G**), were analyzed by flow cytometry for expression of the indicated transcription factor(s). All data are mean ± SEM (p*<0.05, p**<0.005, p***<0.0005, p****<0.00005). (A-G) Representative of 2 experiments.

#### Odc-ΔT mice exhibit defective T_H_17 polarization in an in vivo model of T_H_17 induction

Our *in vitro* experiments highlighted how *Odc*^-/-^ T cells have the capacity to express multiple lineage-defining cytokines and transcription factors within the same cell. However, these data also suggested a requirement for polyamine synthesis in efficient T_H_17 engagement, typified by reduced IL-17 and RORγt expression. Although our T cell transfer colitis experiments supported this conclusion, we tested this further utilizing an *in vivo* model of T_H_17 induction. Treatment with anti-CD3-specific monoclonal antibody drives activation induced cell death in T cells and subsequent engulfment of apoptotic T cell leads to IL-6 and TGF-β production, two cytokines important for the development of T_H_17 cells (Esplugues et al., 2011). These conditions lead to local inflammation in the small intestine and robust T_H_17 cell formation (Esplugues et al., 2011). Indeed, anti-CD3 treatment of WT mice resulted in a large frequency and number of small intestine CD4^+^ T cells expressing IL-17A, IL-17F, and co-expressing these cytokines (**Figs. 2G** and **S3E**). Both the frequency and number of CD4^+^ T cells expressing IL-17A and IL-17A/IL-17F was significantly lower in *Odc*-ΔT mice following anti-CD3 antibody administration (**Figs. 2G** and **S3E**). In this model, more than 70% of small intestine CD4^+^ T cells expressed the T_H_17-defining transcription factor RORγt following anti-CD3 antibody treatment (**Fig. S3F**), however the number and frequency of CD4^+^ T cells expressing RORγt was significantly decreased in *Odc*-ΔT mice (**Figs. 2H** and **S3F**). These data suggest a crucial role for polyamine synthesis in T_H_17 cell formation *in vivo*.

#### Dohh^-/-^ CD4^+^ T cells express dysregulated cytokines and transcription factors across T_H_ cell subsets

Polyamines have pleiotropic roles within mammalian cells (Igarashi and Kashiwagi, 2000). We questioned whether the mechanism through which polyamine synthesis controls T_**H**_ cell subset fidelity is via the production of spermidine, and the use of this critical substrate in the hypusination of eIF5A (**Fig. 3A**). Highlighting the relationship between polyamines and hypusine, eIF5A and hypusinated-eIF5A (eIF5A^H^) increased after T cell activation (**Fig. 3B**), similar to ODC expression (**Fig. 1B**) and polyamines (**Fig. 1C**). Importantly, *Odc*^*-/-*^ T cells exhibited decreased eIF5A^H^, but not total eIF5A (**Fig. 3C**). To test the role of hypusine in CD4^+^ T cells, we crossed mice containing loxP flanked exons 2-4 of *Dohh* (Sievert et al., 2014) with mice expressing *CD4*^*cre*^, to generate mice with *Dohh* specifically deleted in T cells (*Dohh*-ΔT mice). Control mice were absent for cre recombinase. Efficient deletion of DOHH in T cells, and decreased eIF5A^H^, but not total eIF5A, was confirmed by immunoblot (**Fig. 3D**).

**Figure 3.**
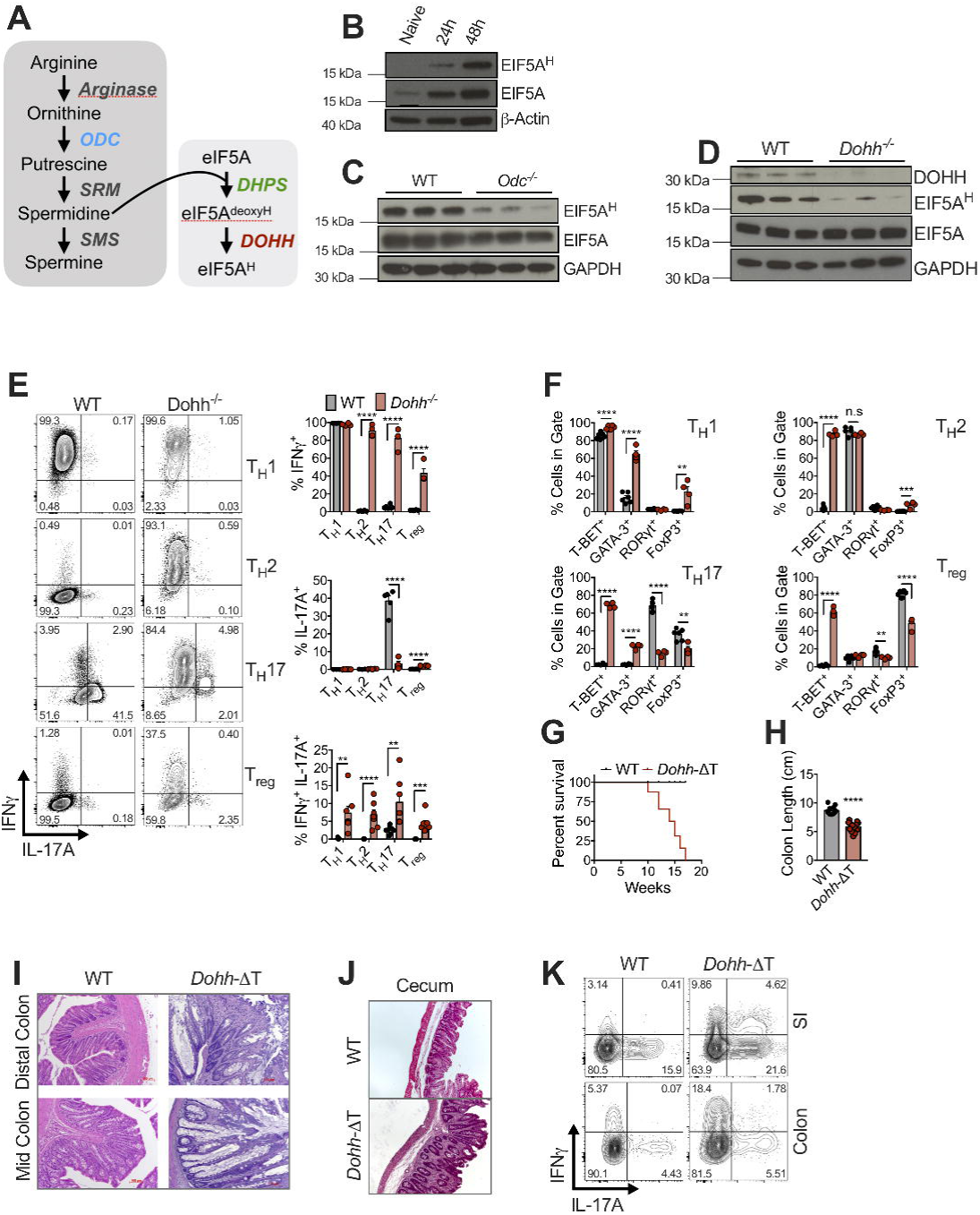
T cell-specific deletion of the hypusine enzyme *Dohh* leads to T cell dysregulation and colitis. (**A**) Schematic of the polyamine biosynthesis pathway and its role in the hypusination of eIF5A. (**B**) Immunoblot analysis of CD4^+^ T cells isolated from spleen of WT mice activated with anti-CD3/CD28 for indicated time or rested overnight in 10 ng/mL IL-7. Representative of 3 biological replicates. (**C**) Immunoblots of WT and *Odc*^-/-^ naïve CD4^+^ T cells activated for 48 hours with anti-CD3/CD28. Representative of 3 biological replicates. N.B. the same loading control and samples were utilized as shown in Fig. 1D. (**D**) Immunoblot of WT and *Dohh*^*-/-*^ naïve CD4^+^ T cells activated with anti-CD3/CD28 for 48 hours. (**E**) Naïve CD4^+^ T cells were isolated from the spleens and lymph nodes of WT and *Dohh*-ΔT mice and polarized for 4 days under indicated T_H_ cell conditions and analyzed for expression of designated cytokine by flow cytometry after 5 hours of restimulation with PMA/ionomycin, representative contour plots are shown. (**F**) Naïve WT and *Dohh*^-/-^ CD4^+^ T were cells activated and polarized as in (E) and analyzed for expression of indicated transcription factor by flow cytometry. Post-natal survival (**G**) and colon length (**H**) of WT and *Dohh*-ΔT mice. Histological images with H&E staining of mid and distal colon (**I**) and cecum (**J**) from 9 week old WT or *Dohh*-ΔT mice. (**K**) CD4^+^ T cells harvested from the indicated organ of 8 week old WT and *Dohh*-ΔT mice were re-stimulated *ex vivo* for 4 hours with PMA/ionomycin and cytokine expression assessed by flow cytometry. (C, D, I, J) Representative of 2 experiments, (E, F) Representative of 4 experiments, (K) Representative of 3 experiments.

We activated naive CD4^+^ T cells from *Dohh*-ΔT mice *in vitro* and polarized them into T_H_1, T_H_2, T_H_17, and T_reg_ cells for 4 days. Upon PMA/ionomycin restimulation, *Dohh*^-/-^ T_H_2, T_H_17, and T_reg_ cells displayed significantly elevated IFN-γ production, with an increased frequency of cells that produced both IFN-γ and IL-17 in all T_H_ cell subsets (**Fig. 3E**). We also detected dysregulated levels of IFN-γ and IL-17 in the supernatant (**Fig. S4A**) as well as increased expression of other cytokines such as TNF, IL-22, and IL-6 (**Fig. S4B**). Furthermore, we observed an increased frequency of cells expressing IL-5, IL-13, and co-expressing these cytokines under T_H_2 conditions (**Fig S4C**), while we also saw an increase in cells co-producing IFN-γ and IL-13 in T_H_2 and T_H_17 culture conditions (**Fig. S4D**). Similar to what we observed for *Odc*^*-/-*^ T cells (**Fig. 1F**), the frequency of T cells producing IL-17 alone decreased in the *Dohh*^*-/-*^ T_H_17 cell culture compared to WT control cells (**Fig. 3E**), while the frequency of T-bet-expressing cells was significantly augmented in all subsets of *Dohh*^*-/-*^ T cells, and the frequency of GATA3 expressing cells was increased in T_H_1 and T_H_17 polarizing conditions (**Figs. 3F** and **S5A**). We also observed a significant reduction in the frequency of cells expressing RORγt under T_H_17 conditions, suggesting a role for DOHH in T_H_17 differentiation (**Figs. 3F** and **S5A)**. The robustness of phenotypic overlap between *Odc*^*-/-*^ and *Dohh*^*-/-*^ T cells suggests that the requirement for polyamine synthesis in enforcing T_H_ cell lineage fidelity is the result of spermidine providing substrate for hypusine synthesis.

#### Mice with a T cell specific deletion of Dohh exhibit T cell dysregulation, inflammation, and colitis

Strikingly, we found that *Dohh*-ΔT mice died at approximately 10-17 weeks of age (**Fig. 3G**). We detected significantly enhanced IFN-γ (**Fig. S5B**) and other cytokines (**Fig. S5C**) in the serum of *Dohh*-ΔT mice compared to control animals, correlating with their disease. We observed decreased colon length in *Dohh*-ΔT mice (**Fig. 3H**) and histology revealed increased immune infiltrates and altered villi structure in the colons of *Dohh*-ΔT mice (**Fig. 3I**) and caecal thickening (**Fig. 3J**), indicative of colitis, a T cell-driven pathology (Powrie et al., 1994). T cells isolated from the small intestine and colon (**Figs. 3K** and **S5D**), as well as from the lungs and spleen (**Fig. S5D**), of *Dohh*-ΔT mice displayed increased frequencies of CD4^+^ T cells producing IFN-γ, and IFN-γ and/or IL-17, upon restimulation *ex vivo*, directly correlating their intestinal inflammation with an increase in cytokine producing T cells *in vivo* (Harbour et al., 2015; Park et al., 2005; Wang et al., 2015; Zielinski et al., 2012). CD4^+^ T cells isolated from the colon or spleen of *Dohh*-ΔT mice also displayed increased T-bet expression (**Fig. S6A**), without any difference observed in the frequency of Foxp3 expressing T cells in these tissue sites (**Fig. S6B**). These data suggest an increase in inflammatory T_H_ cells, rather than a loss of T_reg_ cells, underlies the observed pathology in *Dohh*-ΔT mice.

#### T cell specific deletion of Dhps leads to T cell dysregulation, inflammation, and colitis

The similarity of the phenotypes between *Odc*^*-/-*^ and *Dohh*^*-/-*^ T cells indicated that the role of polyamine synthesis in T_H_ cell subset specification was mechanistically linked to hypusine synthesis. We reasoned that if this were true, we should observe a similar phenotype to *Odc*^*-/-*^ and *Dohh*^*-/-*^ T cells in a third genetic model, i.e. *Dhps* deficiency (**Fig. 3A**). We crossed mice containing loxP flanked exons 2-7 of the *Dhps* gene (Pällmann et al., 2015) with mice expressing *CD4*^*cre*^, to generate mice deleted for *Dhps* in T cells (*Dhps*-ΔT mice). Control mice were absent for cre recombinase. Efficient deletion of DHPS in T cells, and decreased eIF5A^H^, but not total eIF5A, was confirmed by immunoblot (**Fig. 4A**). Like *Dohh*^*-/-*^ T cells, *in vitro* activated CD4^+^ T cells from *Dhps*-ΔT mice exhibited significantly elevated IFN-γ, with increased frequencies of cells producing both IFN-γ and IL-17 (**Fig. 4B**), increased T-bet expression in T_H_2, T_H_17, and T_reg_ cells (**Fig. S7**), and decreased frequencies of cells producing RORγt and IL-17 alone under T_H_17 conditions (**Figs. 4B** and **S7**). As with *Dohh*-ΔT mice, two independently generated lines of *Dhps*-ΔT mice in distinct animal facilities, developed fatal disease at around 10-20 weeks of age (**Fig. 4C**) with intestinal inflammation as measured by significantly decreased colon length (**Fig. 4D**). *Dhps*-ΔT mice also displayed increased frequencies of IFN-γ, and IFN-γ and/or IL-17 producing CD4^+^ T cells in the lung, colon, and spleen (**Fig. S8A)**, increased T-bet and RORγt expression **(Fig. S8B)**, correlating with their inflammatory phenotype (Harbour et al., 2015; Neurath et al., 2002; Yen et al., 2006), with no change in the frequency of Foxp3 expressing cells (**Fig. S8C**).

**Figure 4.**
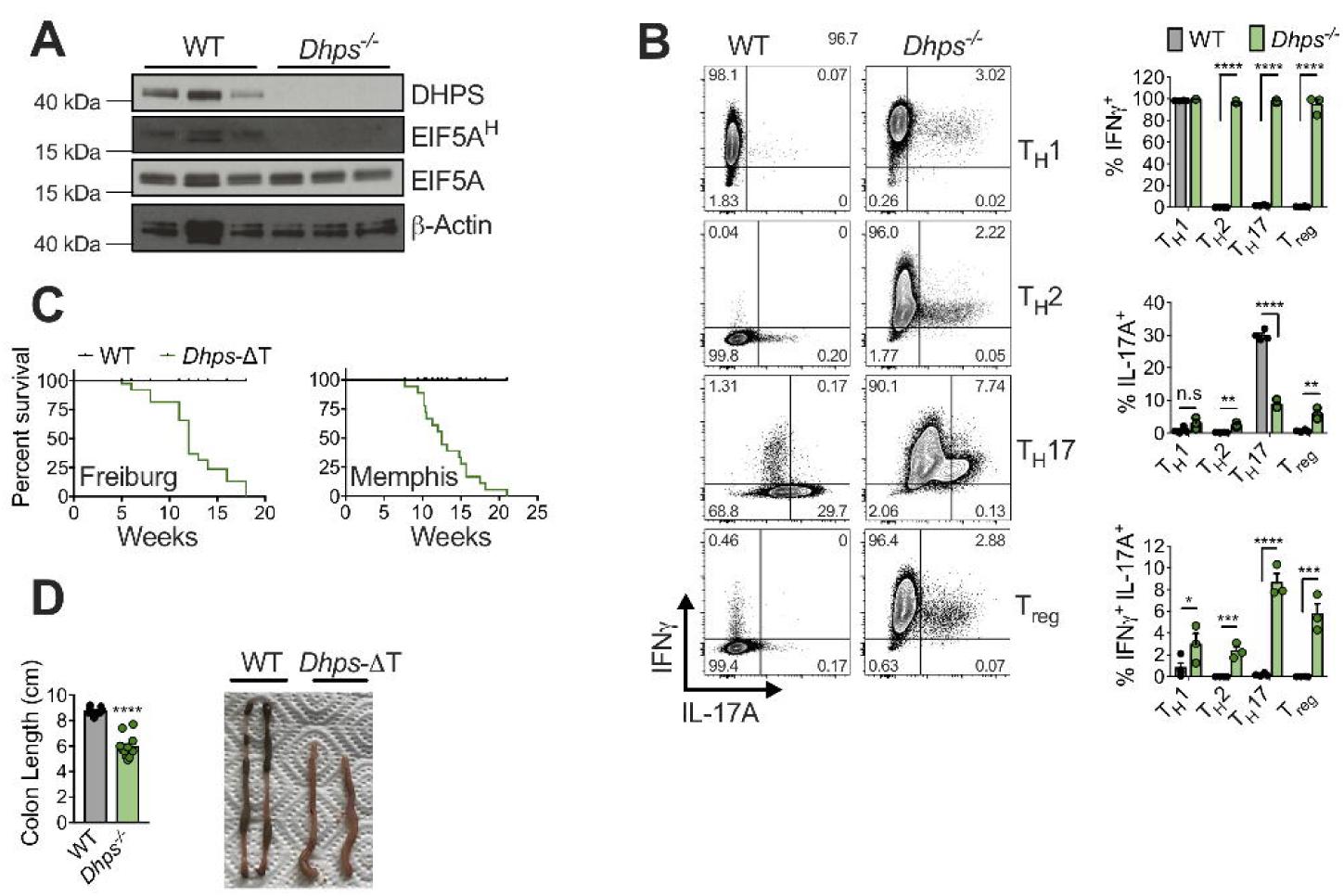
T cell-specific deletion of *Dhps* confirms a role for hypusine synthesis in T_H_ lineage fidelity. (**A**) Immunoblot of WT and *Dhps*^*-/-*^ CD4^+^ T cells activated for 48 hours with anti-CD3/CD28. (**B**) WT and *Dhps*^-/-^ naïve CD4^+^ T cells were activated with anti-CD3/CD28 for 4 days in indicated T_H_ cell polarizing conditions. Cytokine expression was assessed by flow cytometry 5 hours after PMA/ionomycin re-stimulation, representative contour plots are shown. (**C**) Post-natal survival of WT and *Dhps*-ΔT mice from Freiburg or Memphis facilities. (**D**) Colon length and representative images from 6 week old WT and *Dhps*-ΔT mice. All data are mean ± SEM (p*<0.05, p**<0.005, p***<0.0005, p****<0.00005). (A) Representative of 1 experiment, (B) Representative of 3 experiments.

#### Polyamine or hypusine-deficient CD4^+^ T cells exhibit widely altered chromatin accessibility linked to broad changes in histone methylation and acetylation

The strikingly similar phenotype observed in *Odc-, Dohh-*, and *Dhps*-deficient T cells highlighted the strong mechanistic association between polyamine synthesis and hypusine in regulating T_H_ cell subset fidelity. Given that *Odc-, Dohh-*, and *Dhps*-deficient T cells displayed attributes of multiple T_H_ cell subsets at once, indicating a lack of focused T_H_ cell lineage commitment, we performed ATAC-Seq across the genotypes and T_H_ cell subsets to assess chromatin accessibility. Chromatin accessibility influences the access of T_H_ cell subset-specific transcription factors that drive expression of T_H_ cell subset-specific genes (Hirahara et al., 2011). PCA analysis revealed that chromatin accessibility was highly disparate between all *Odc*^*-/-*^ and *Dohh*^*-/-*^ T_H_ cell subsets when compared to WT cells (**Fig. 5A**). Across T_H_ cell subsets, many differentially regulated regions of accessible chromatin were shared between *Odc*^*-/-*^ and *Dohh*^*-/-*^ T_H_ cells (**Fig. 5B**), including at loci critical for T_H_ cell differentiation and lineage identity, such as *Tbx21* (T-Bet), *Gata3, Rorc, Ifng*, and *Il17* (**Fig. S9A**). In parallel, we also performed RNA-Seq to assess gene transcription. PCA analysis suggested a significantly altered transcriptional profile in both *Odc*^*-/-*^ and *Dohh*^*-/-*^ cells relative to WT cells across all T_H_ cell subsets (**Fig. 5C**) with many genes found to be commonly differentially regulated in both genotypes (**Fig. 5D**), including genes essential for T_H_ cell differentiation and effector function (**Fig S9B**).

**Figure 5.**
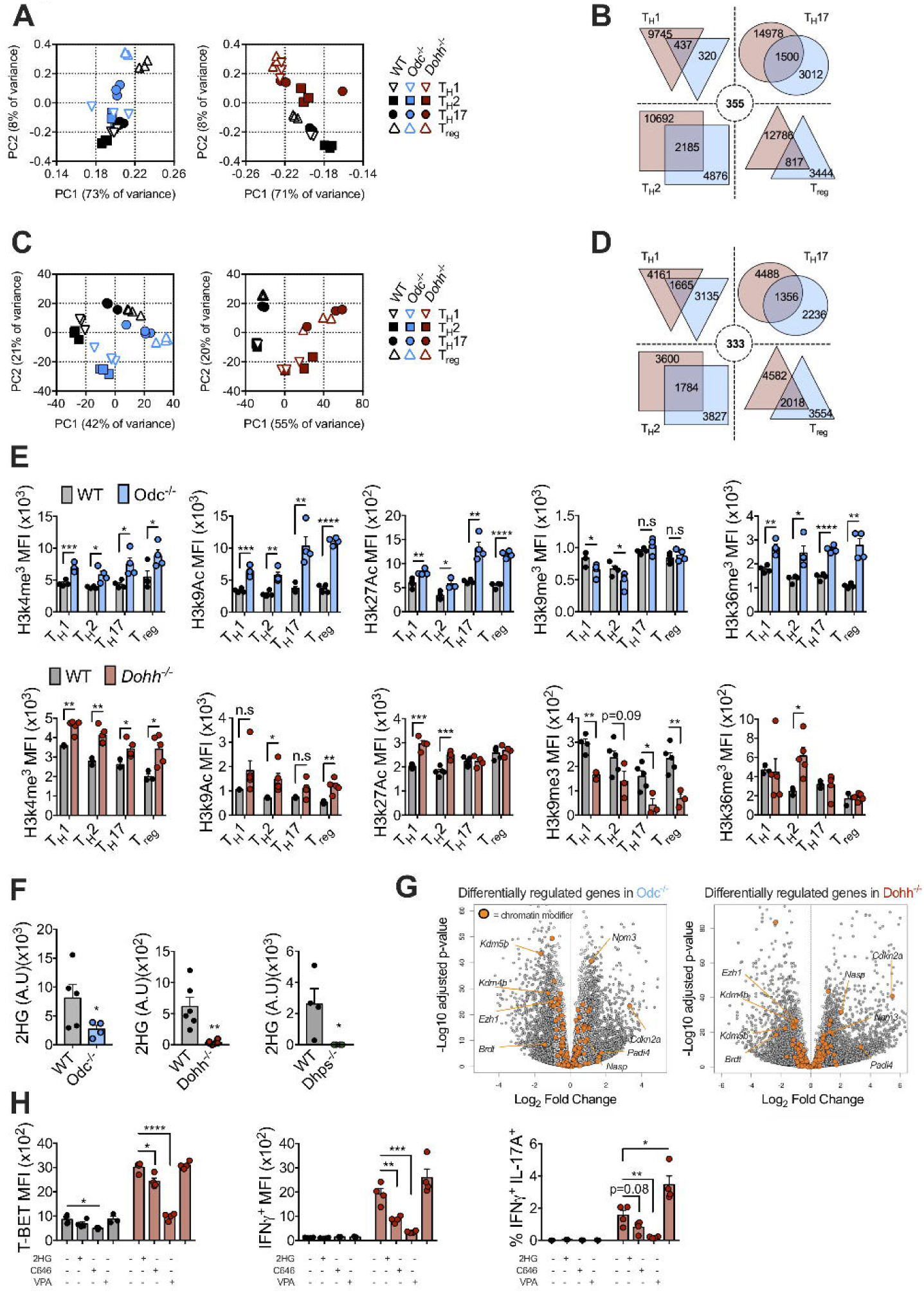
The polyamine-hypusine axis regulates the T cell epigenome to enforce appropriate T_H_ cell differentiation and function. (**A**) ATAC-Seq was performed on naïve CD4^+^ T cells isolated from the spleens and lymph nodes of WT, *Odc*-ΔT, and *Dohh*-ΔT mice and activated for 4 days in indicated T_H_ cell polarizing conditions. The reproducibility of each treatment and the effect of *Odc* and *Dohh* deficiency are shown via principal component analysis, indicating the percentage of variance allocated to each component in parenthesis. (**B**) Venn diagram depicting the number of differentially regulated regions of open chromatin that are disparate between, or shared between, *Odc*^*-/-*^ and *Dohh*^*-/-*^ CD4^+^ T cells of the indicated T_H_ cell lineage. Dashed circle indicates number of differentially regulated regions of open chromatin that are common among all T_H_ cell subsets. (**C**) RNA was obtained and sequenced from naïve CD4^+^ T cells isolated from the spleens and lymph nodes of WT, *Odc*-ΔT and *Dohh*-ΔT mice and activated for 4 days under indicated T_H_ cell polarizing conditions. The reproducibility of each treatment and the effect of *Odc* and *Dohh* deficiency are shown via principal component analysis, indicating the percentage of variance allocated to each component in parenthesis. (**D**) Venn diagram depicting the number of differentially regulated genes disparate between, or shared between, *Odc*^*-/-*^ and *Dohh*^*-/-*^ CD4^+^ T cells of the indicated T_H_ lineage. Dashed circle indicates number of differentially expressed genes that are common among all T_H_ cell subsets. (**E**) WT, *Odc*^*-/-*^ or *Dohh*^*-/-*^ CD4^+^ T cells assayed for chromatin modifications by flow cytometry in indicated T_H_ cell subset 4 days post-activation/polarization. Analysis was performed on Ki-67^+^ cells and diploid cells with ‘single’ DNA content based on FxCycle (dapi) staining in the live cell gate. (**F**) Naïve CD4^+^ T cells from *Odc*-ΔT, *Dohh*-ΔT, and *Dhps*-ΔT were activated with anti-CD3/CD28 for 48 hours and the quantity of 2-hydroxyglutarate was assessed by GC-MS. (**G**) Volcano plot depicting differentially regulated genes present in all T_H_ subsets in *Odc*^*-/-*^ and *Dohh*^*-/-*^ CD4^+^ T cells relative to WT cells. Dark grey circles represent statistically significant changes in expression; orange circles represent genes known to act as chromatin modifiers; annotated chromatin modifiers are differentially expressed in both *Odc*^*-/-*^ and *Dohh*^*-/-*^ across all T_H_ subsets. (**H**) Naïve CD4^+^ T cells from WT and *Dohh*-ΔT mice were activated and polarized under T_H_2 conditions. After 24 hours cells were treated with either 25 mM D-2-HG, 20 μM C646 (HAT inhibitor), or 400 μM valproic acid (VPA, HDAC inhibitor) and the expression of indicated protein was assessed 48 hours later. All data are mean ± SEM (p*<0.05, p**<0.005, p***<0.0005, p****<0.00005). (E) Representative of 1-3 experiments, (F) Representative of 2 experiments, (H) Representative of 3 experiments.

Since we observed vast differences in chromatin accessibility, we measured epigenetic marks on histone proteins, including activating modifications such as H3k4me^3^, H3k9Ac, H3k27Ac, and H3k36me^3^, as well as the repressive mark H3k9me^3^ (Bannister and Kouzarides, 2011; Lawrence et al., 2016) between *Odc*^*-/-*^, *Dohh*^*-/-*^, and control T_H_ cell subsets. We found significant increases in many marks that correlate with transcriptional activation on histones across all *Odc*^*-/-*^ and *Dohh*^*-/-*^ T_H_ cell subsets, with a decrease in H3k9me^3^, which correlates with transcriptional repression, in several T_H_ cell subsets when compared to control WT cells (**Fig. 5E**). These data show that *Odc*^*-/-*^ and *Dohh*^*-/-*^ T cells had highly remodeled chromatin, and indicated a prominent role of the polyamine-hypusine axis in regulating the T cell epigenome.

#### Polyamine or hypusine-deficient CD4^+^ T cells have altered TCA cycle metabolites with low 2-HG

Alterations in metabolites, such as those in one-carbon metabolism, which directly provide carbon substrate for methylation reactions (Serefidou et al., 2019), or those in the TCA cycle, such as α-ketoglutarate (α-KG), which affect the catalytic activity of α-KG-dependent dioxygenases, including the Tet lysine demethylase proteins (Lu and Thompson, 2012), profoundly influence histone modifications. To ascertain how the marked differences in the epigenetic landscape (**Fig. 5E**) of *Odc*^*-/-*^ and *Dohh*^*-/-*^ T_H_ cell subsets could manifest, we next performed a metabolomics analysis on *Odc*^*-/-*^, *Dohh*^*-/-*^, and *Dhps*^*-/-*^ naïve CD4^+^ T cells activated for 48 hours with anti-CD3/CD28 and IL-2. We found a general decrease in TCA cycle metabolites, including α-KG (**Fig. S10A**), with a significant loss of 2-hydroxyglutarate (2-HG) when compared to control cells (**Fig. 5F**). 2-HG is produced from α-KG, sometimes in settings of dampened mitochondrial activity (Lu and Thompson, 2012), and can function as an antagonist of α-KG-dependent enzymes to modulate histone methylation and chromatin accessibility (Chowdhury et al., 2011; Lu et al., 2012; Xu et al., 2011). Decreased TCA cycle metabolites were also evident in our prior work showing that the hypusine pathway is critically important for mitochondrial activity and TCA cycle integrity in MEFs and bone marrow derived macrophages (Puleston et al., 2019).

In addition to alterations in the metabolic status of cells influencing chromatin accessibility and gene expression, we also considered that expression of chromatin-modifying enzymes themselves could be dysregulated and contribute to the phenotype in polyamine-hypusine deficient T cells. Analysis of our RNA-Seq data showed significant changes in the expression of chromatin modifiers, many of which were shared between *Odc*^*-/-*^ and *Dohh*^*-/-*^ across all T_H_ cell subsets (**Fig. 5G**). Taken together, these data prompted us to postulate that changes in the TCA cycle intermediate 2-HG combined with differential expression of chromatin modifiers, could influence the epigenetic state of polyamine-hypusine deficient T_H_ cells and contribute to their dysregulation and lack of focused lineage commitment.

#### 2-HG or HAT inhibition restore normal cytokine and transcription factor expression in polyamine or hypusine-deficient CD4^+^ T cells

Thus, we reasoned that either deceased levels of 2-HG, or changes in the expression of chromatin modifiers, or both, could alter histone methylation and/or acetylation and lead to the promiscuous expression of cytokines and transcription factors across T_H_ cell subsets in polyamine-hypusine deficient T cells. We tested whether restoring 2-HG or decreasing histone acetylation could correct inappropriate lineage commitment. We cultured *Odc*^*-/-*^ and *Dohh*^*-/-*^ T_H_ cells under T_H_2 polarizing conditions in the presence of a histone acetyltransferase (HAT) inhibitor C646 (Bowers et al., 2010), or 2-HG and assessed expression of IFN-γ, IL-17, and T-bet. We chose T_H_2 conditions because WT T_H_2 cells should not express any of these proteins to a great degree, however these proteins were significantly dysregulated in polyamine-hypusine deficient T cells (**Figs. 1**-**4**). We found the frequency of IFN-γ, IFN-γ and IL-17, and T-bet expressing *Odc*^*-/-*^ and *Dohh*^*-/-*^ T_H_2 cells were restored to almost WT levels after C646 or 2-HG treatment (**Figs. 5H** and **S10B**), while GATA-3 expression was unaffected by these treatments (**Fig. S10C**). Of note, valproic acid (VPA), a histone deacetylase (HDAC) inhibitor, either had no effect or increased the expression of these proteins (**Figs. 5H, S10B**, and **S10C**). These results suggest that linked alterations to metabolism and chromatin accessibility contribute to the transcriptional dysregulation evident in polyamine-hypusine deficient CD4^+^ T_H_ cells.

## DISCUSSION

Polyamines are involved in many processes, including gene transcription and translation, chromatin structure, cell growth and proliferation, aging, autophagy, and ion channel function, but a complete mechanistic understanding of how they exert their biological functions is often lacking. Here we provide genetic evidence linking polyamine synthesis to the hypusine enzymes DHPS and DOHH in activated CD4^+^ T_H_ cells and demonstrate that this axis is critical for maintaining the epigenome to focus T_H_ cell subset fidelity.

In addition to synthesizing polyamines, cells can acquire them from the extracellular environment (Abdulhussein and Wallace, 2014). Cells utilize polyamines derived from the diet, as well as those from the microbiota, but how the transport of these molecules is achieved and regulated across cell types is unclear. Intracellular polyamines must be tightly controlled, as too high concentrations can be toxic, and too low inhibits cell proliferation (Pegg, 2016). Requirements for polyamines will change with cell activity. It was interesting that colitis manifested so strongly in the hypusine-deficient mice, but was only conferred by T cell transfer in the ODC-deficient setting. These data suggest that T cells in the gut have a higher requirement for hypusine upon adoptive transfer in the colitis model, than in the steady state. It stands to reason that *Odc*^*-/-*^ T cells can still acquire some polyamines from the diet and/or microbes and use these for hypusine synthesis, whereas *Dohh*- and *Dhps*-deficient T cells cannot synthesize hypusine, even if polyamine substrates are present. Our data showing that putrescine rescues cytokine dysregulation in *Odc*^*-/-*^ T cells supports this idea. These data have implications for immune mediated diseases, where T cells have critical roles in maintaining tolerance to self-antigens and tissue homeostasis. Future research will focus on the contribution of diet and microbe-derived polyamines *in vivo* versus polyamine synthesis.

While polyamines are known to influence cell proliferation, exactly how these molecules influence this process is not well defined. *Odc*^*-/-*^, *Dohh*^-/-^, and *Dhps*^-/-^ T cells proliferate less than WT T cells *in vitro*, and *in vivo* (data not shown), and *Dohh*-ΔT and *Dhps*-ΔT mice have lower numbers of peripheral T cells, although different T cell types are present at frequencies similar to control mice (data not shown). Whether dampened proliferation is required for the dysregulated cytokines and active chromatin state of these cells is not known at this time. In T cells, it may not be possible to tease apart proliferation and differentiation, as these processes are linked in these cells (Jelley-Gibbs et al., 2000). However, it has been shown that deletion of *Odc* in macrophages causes increased inflammation due to alterations in histone modifications, resulting in changes in chromatin structure and up-regulated transcription in these cells (Hardbower et al., 2017). Whether or not hypusine plays a role in this process in macrophages has yet to be investigated, but these data suggest that polyamines can influence the epigenome in cells regardless of cell division, as inflammatory macrophages are a non-proliferating cell type.

We previously investigated the polyamine-hypusine pathway in MEFs and bone marrow derived macrophages and found that it was critically important for TCA cycle integrity by maintaining the efficient expression of a subset of mitochondrial and TCA cycle enzymes. In these studies we identified a loss of TCA cycle metabolites after acute polyamine depletion or DHPS inhibition (Puleston et al., 2019). In our current study, using models of chronic gene deletion of *Odc, Dohh*, and *Dhps* in T cells, we observed a similar perturbation in core TCA cycle metabolites. Overall, a picture emerges that polyamines and hypusine have specific effects on the mitochondria, but determining how this regulation occurs requires further study. The only known functions of DOHH and DHPS are to, in a two-step process, hypusinate eIF5A (Park et al., 2010). Based on this it is logical that changes in eIF5A function due to a failure of hypusination must be critical for the changes in CD4^+^ T cell function reported here. Functionally, eIF5A is a translation factor that when hypusinated preferentially regulates the translation of transcripts with specific sequence properties (Gutierrez et al., 2013; Pelechano and Alepuz, 2017; Schuller et al., 2017). What this subset of transcripts is in differentiating CD4^+^ T cells, and how their translation might impact the chromatin and transcriptional states of these cells, remains to be determined. The alternative hypotheses, that hypusination is critical for other events unrelated to eIF5A, or that DHPS and DOHH have additional functions unrelated to hypusination, remain possible, although given the current state of the field, unlikely. What is clear, however, is that DHPS and DOHH have profound effects on CD4^+^ T cell differentiation and subset fidelity.

Chromatin accessibility facilitates the access of transcription factors to DNA and is an important underlying component of identity specification in T_H_ cell subsets (Murphy and Stockinger, 2010). Epigenetic marks such as those on histones can modulate and/or reinforce chromatin states and influence gene expression (Allis and Jenuwein, 2016). Recent reports have demonstrated a crucial role for 2-HG in regulating the epigenetic state of T cells and how this impacts T cell differentiation and function (Weinberg et al., 2019; Xu et al., 2017), and our data point to a loss of 2-HG as a factor in the epigenetic dysregulation observed in *Odc* and *Dohh* deficient T cells. 2-HG regulates chromatin by acting as an antagonist of TET demethylases (as well as other α-KG-dependent dioxygenases). It is produced from the TCA cycle intermediate α-KG by malate dehydrogenase or lactate dehydrogenase under specific conditions related to hypoxia or changes in mitochondrial electron transport chain activity (Weinberg et al., 2019). The fact that either exogenous 2-HG, or HAT inhibition, rescues the aberrant cytokine and transcription factor expression in *Odc* and *Dohh* deficient T cells demonstrates that a major feature of the polyamine/hypusine pathway in CD4^+^ T cell differentiation is to modify and maintain the epigenome for appropriate lineage commitment.

Our results showing that purified naïve *Odc*^*-/-*^ CD4^+^ T cells are more colitogenic than control WT cells in a T cell transfer model of IBD illustrate that a defect in this pathway alone in naïve CD4^+^ T cells can drive disease. Although we do not see altered frequencies of Foxp3^+^ T cells in this setting, we do see dysregulated IFN-γ production in *Odc-, Dohh-*, and *Dhps*-deficient Foxp3^+^ T cells *in vitro*, suggesting that dysregulation in T_reg_ function could synergize with exaggerated inflammatory effector T_H_ cell functions to precipitate the development of colitis.

CD4-cre deletes in all cells that express *Cd4* during development, and therefore CD8^+^ T cells were also deficient in the polyamine-hypusine axis in our studies, and in initial analyses these cells also exhibited extensive gene expression dysregulation (data not shown). The contribution of effects in these cells to the disease states described herein remains to be determined. However, the colitis observed here in *Dohh*-ΔT and *Dhps*-ΔT mice is believed to be primarily a CD4^+^ T cell-mediated disease (Shale et al., 2013), and in the *Odc*^*-/-*^ T cell transfer model accelerated development of colitis was mediated by purified CD4^+^ T cells alone.

Determining why IL-17 production is dampened within the T_H_17 lineage upon *Odc-, Dohh-*, or *Dhps-*deletion requires further study. However, these results illustrate that depending on context, the polyamine-hypusine axis induces the repression of some cytokine genes. Further analysis of the relatedness of genes that are repressed in polyamine-hypusine deficient CD4^+^ T cells compared to those that are expressed more strongly may allow description of shared features that determine outcome. In future studies it will be informative to establish how the polyamine-hypusine pathway influences a variety of immune cells and how this contributes to health and disease.

## FIGURE LEGENDS

**Figure S1.**
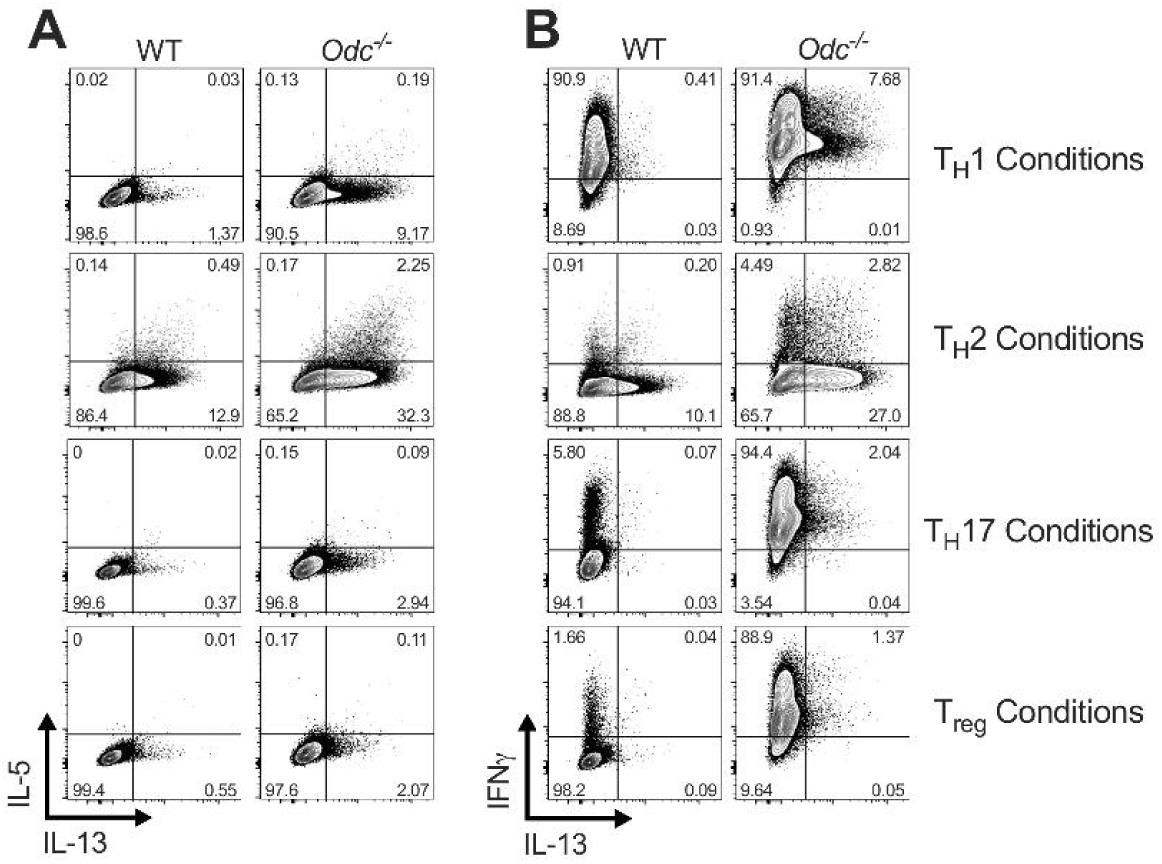
Cytokine expression in *in vitro* activated *Odc*^*-/-*^ CD4^+^ T cells (Related to Figure 1). (**A** and **B**) WT and *Odc*^-/-^ naïve CD4^+^ T cells were activated with anti-CD3/CD28 for 4 days in indicated T_H_ cell polarizing conditions. Representative flow cytometry plots of intracellular cytokine expression 5 hours after PMA/ionomycin re-stimulation are shown, representative of 5 experiments.

**Figure S2.**
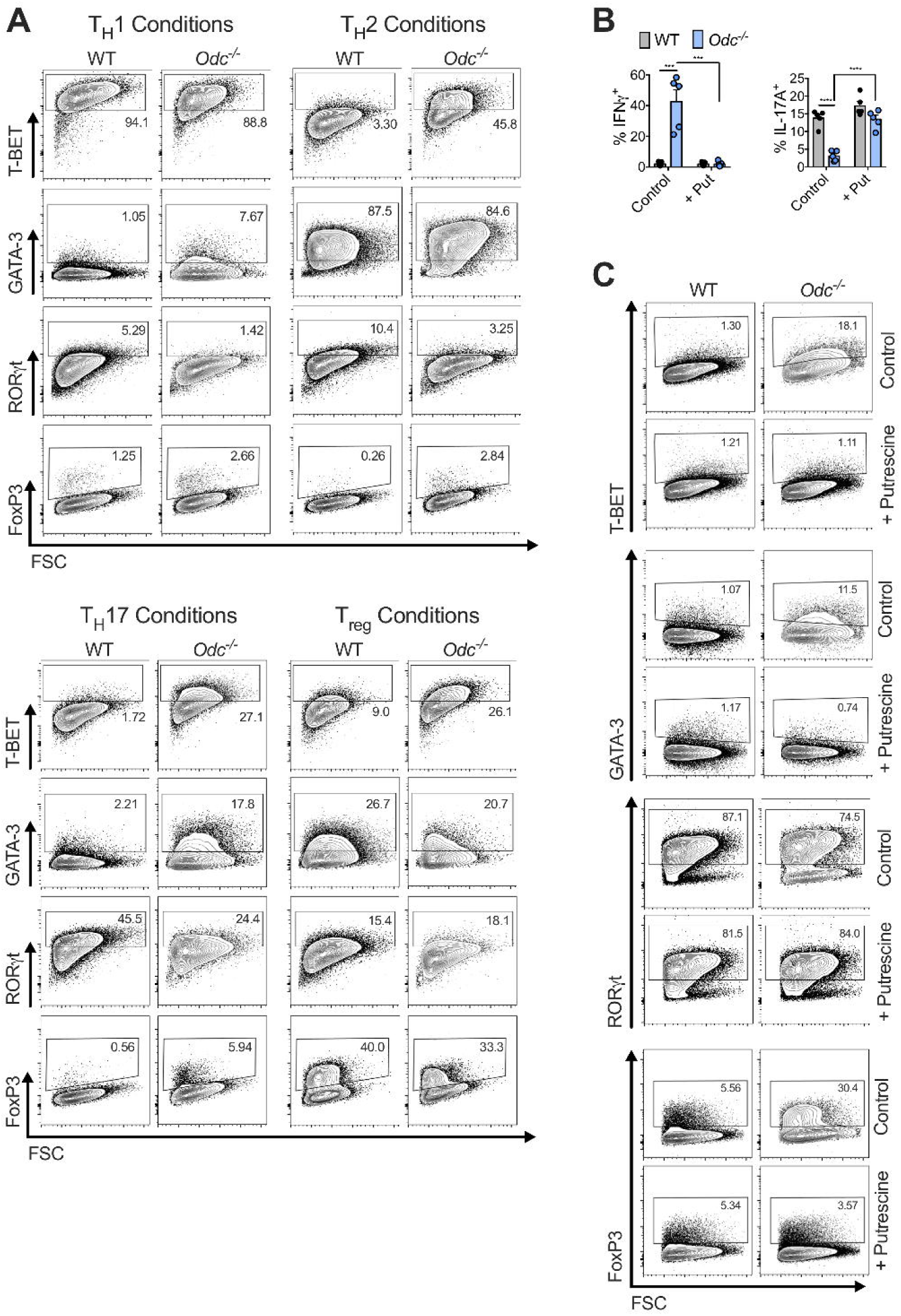
Transcription factor expression in, and putrescine treatment of, *in vitro* activated *Odc*^*-/-*^ CD4^+^ T cells (Related to Figure 1). (**A**) Representative flow cytometry plots showing expression of indicated transcription factors in WT and *Odc*^-/-^ naïve CD4^+^ T cells activated and polarized under the designated T_H_ cell conditions. (**B**) WT and *Odc*^-/-^ naïve CD4^+^ T cells activated under T_H_17 cell polarizing conditions ± 250 μM putrescine for 4 days. Intracellular cytokine was assessed 5 hours post re-stimulation with PMA/ionomycin. (**C**) WT and *Odc*^-/-^ CD4^+^ T cells treated as in (B), representative contour plots with expression of labeled transcription factors are shown. All data are mean ± SEM (p*<0.05, p**<0.005, p***<0.0005, p****<0.00005). Representative of 5 experiments, (B, C) Representative of 2 experiments.

**Figure S3.**
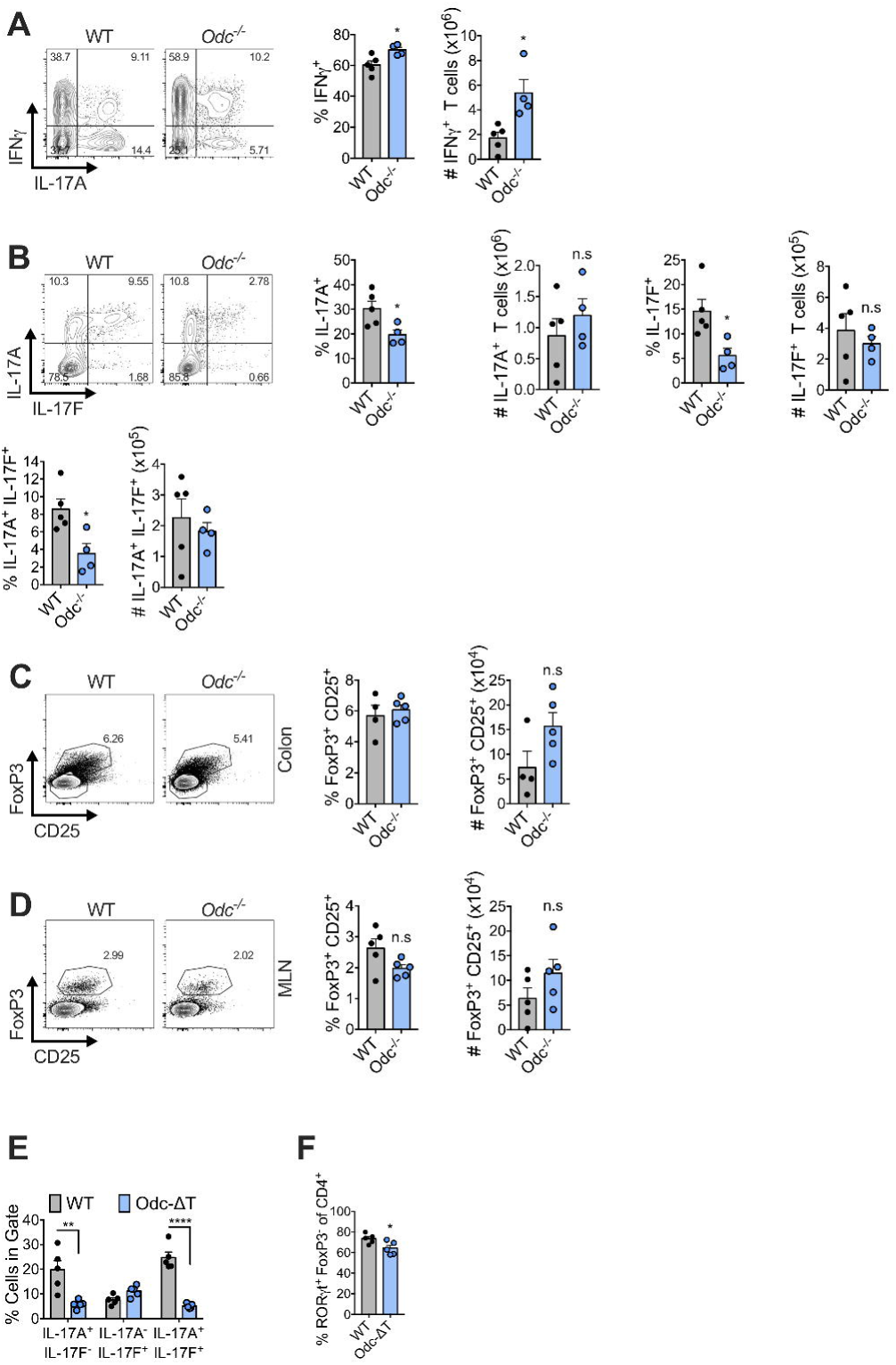
Cytokine and transcription factor expression in *Odc*^*-/-*^ CD4^+^ T cells isolated *ex vivo* (Related to Figure 2). (**A** and **B**) 4×10^5^ WT or *Odc*^*-/-*^ naïve (CD45RB^hi^ CD25^-^ CD44^lo^ CD62L^hi^) CD4^+^ T cells were adoptively transferred into *Rag1*^*-/-*^ recipient mice. On day 37, the frequency and number of CD4^+^ T cells in the MLN expressing the indicated cytokine was analyzed following 4 hours *ex vivo* re-stimulation with PMA/ionomycin. Frequency and number of T_regs_ in colon (**C**) and MLN (**D**) (day 37 post-transfer) from *Rag1*^*-/-*^ mice that received either WT or *Odc*^*-/-*^ CD4^+^ T cells. (**E** and **F**) WT and *Odc*-ΔT mice were treated 50 μg anti-CD3 antibody by intraperitoneal injection on day 0, day 2, and day 4 and sacrificed 4 hours after the last injection. The frequency of CD4^+^ T cells expressing indicated cytokine (**E**) and transcription factor (**F**) was assessed from the small intestine by flow cytometry. All data are mean ± SEM (p*<0.05, p**<0.005, p***<0.0005, p****<0.00005). (A-F) Representative of 2 experiments.

**Figure S4.**
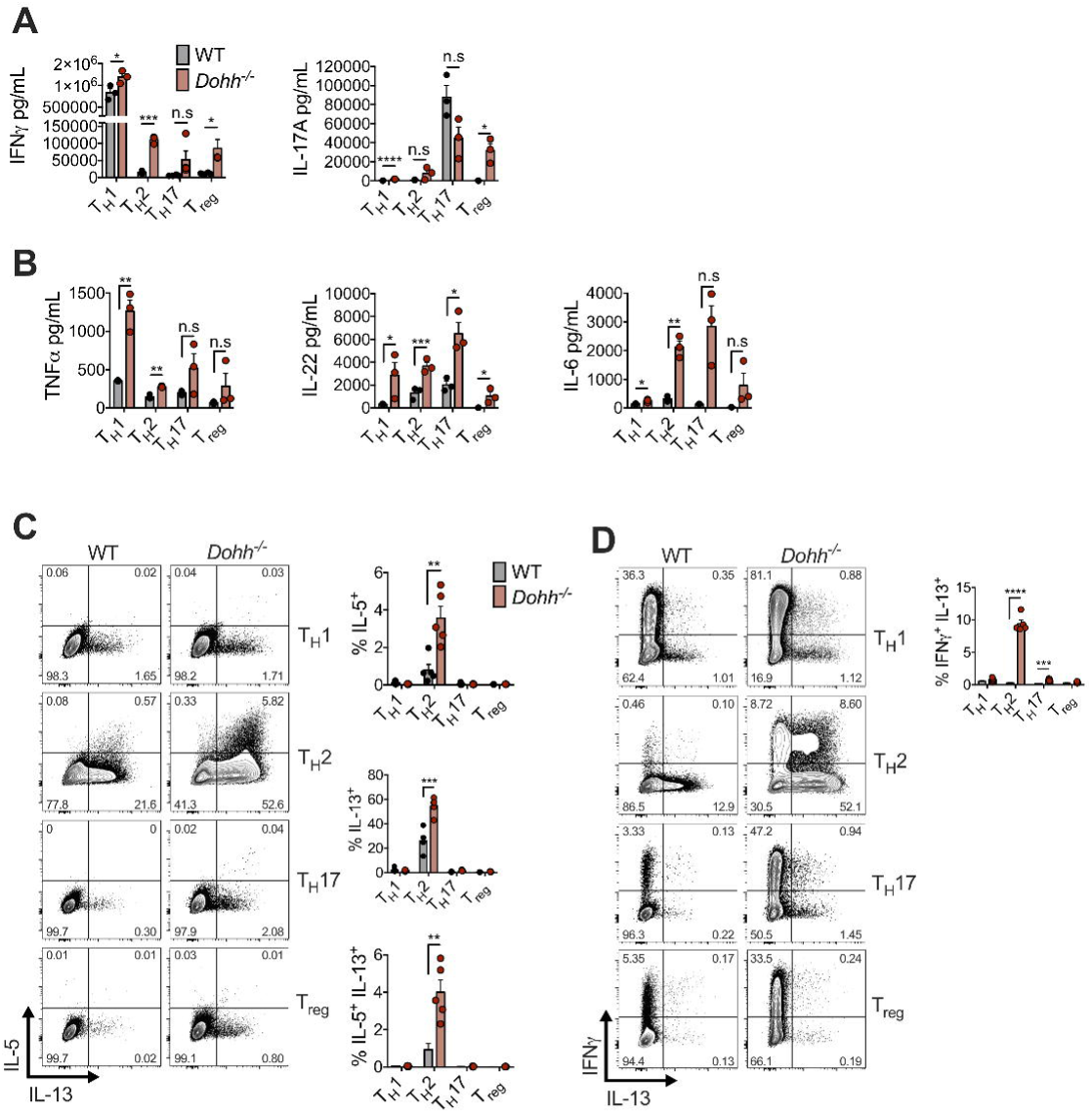
Cytokine expression by *in vitro* activated *Dohh*^*-/-*^ CD4^+^ T cells (Related to Figure 3). (**A** and **B**) WT and *Dohh*^-/-^ naïve CD4^+^ T cells were activated for 4 days under T_H_ cell polarizing conditions. Culture supernatent was analyzed for presence of indicated cytokine by cytokine bead array. (**C** and **D**) WT and *Dohh*^-/-^ naïve CD4^+^ T cells were polarized as in (A) and assayed for the expression of designated cytokine by flow cytometry after 5 hours of PMA/ionomycin re-stimulation. Representative contour plots are also shown. All data are mean ± SEM (p*<0.05, p**<0.005, p***<0.0005, p****<0.00005). (A-D) Representative of 2 experiments.

**Figure S5.**
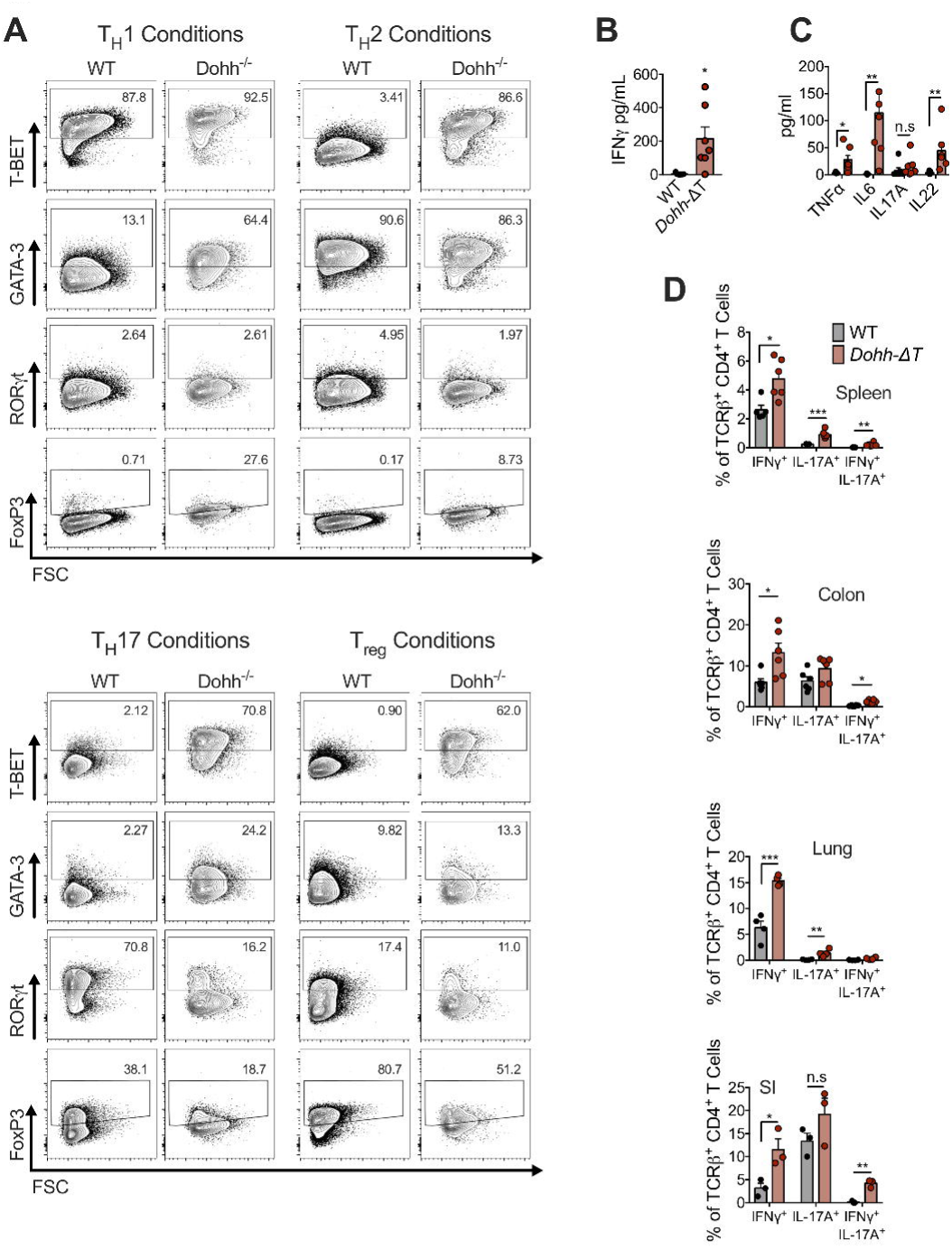
Transcription factor and cytokine expression by *Dohh*^*-/-*^ CD4^+^ T cells and *Dohh*-ΔT mice. (Related to Figure 3). (**A**) WT and *Dohh*^-/-^ naïve CD4^+^ T cells were polarized for 4 days under T_H_ cell conditions and the expression of indicated transcription factor assessed by flow cytometry. Representative contour plots are shown. Serum from 6-10 week old WT and *Dohh*-ΔT mice was assayed for IFN-γ (**B**) and other indicated cytokines (**C**) by cytokine bead array. (**D**) CD4^+^ T cells were harvested from designated organs of WT and *Dohh*-ΔT and cytokine expression was analyzed by flow cytometry following re-stimulated for 4 hours with PMA/ionomycin (SI = small intestine). All data are mean ± SEM (p*<0.05, p**<0.005, p***<0.0005, p****<0.00005).(A) Representative of 4 experiments, (B-D) Representative of 3 experiments.

**Figure S6.**
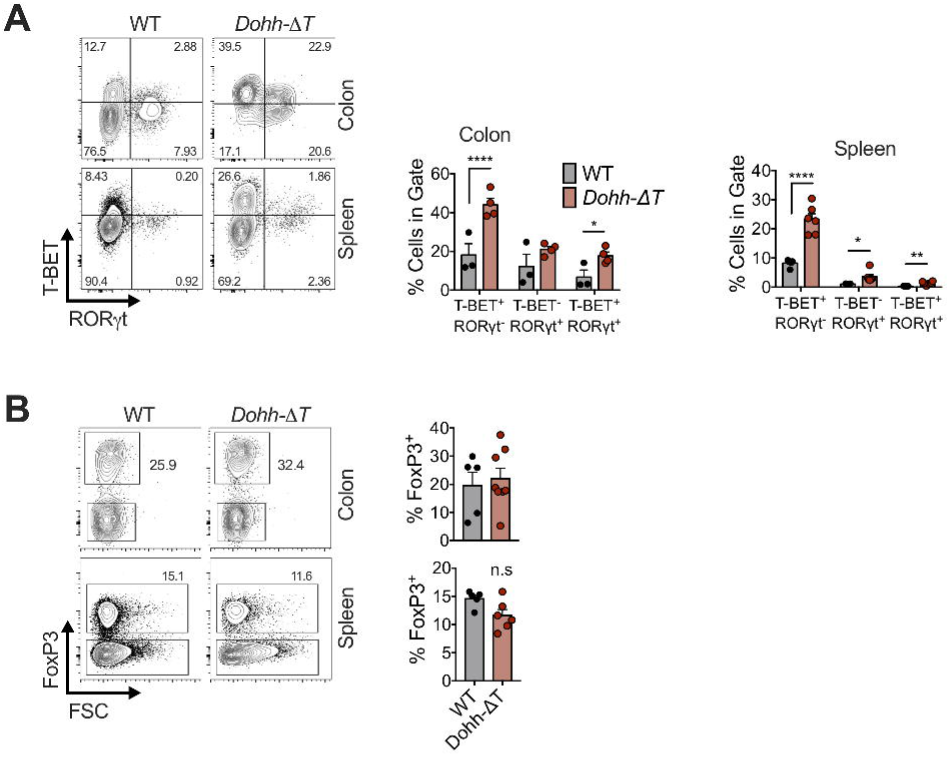
Transcription factor expression of *Dohh*^*-/-*^ CD4^+^ T cells isolated *ex vivo* (Related to Figure 3). (**A** and **B**) CD4^+^ T cells were harvested from designated organs of WT and *Dohh*-ΔT and expression of indicated transcription factor was analyzed by flow cytometry. Representative contour plots are shown and gated on CD4^+^ T cells. All data are mean ± SEM (p*<0.05, p**<0.005, p***<0.0005, p****<0.00005), representative of 3 experiments.

**Figure S7.**
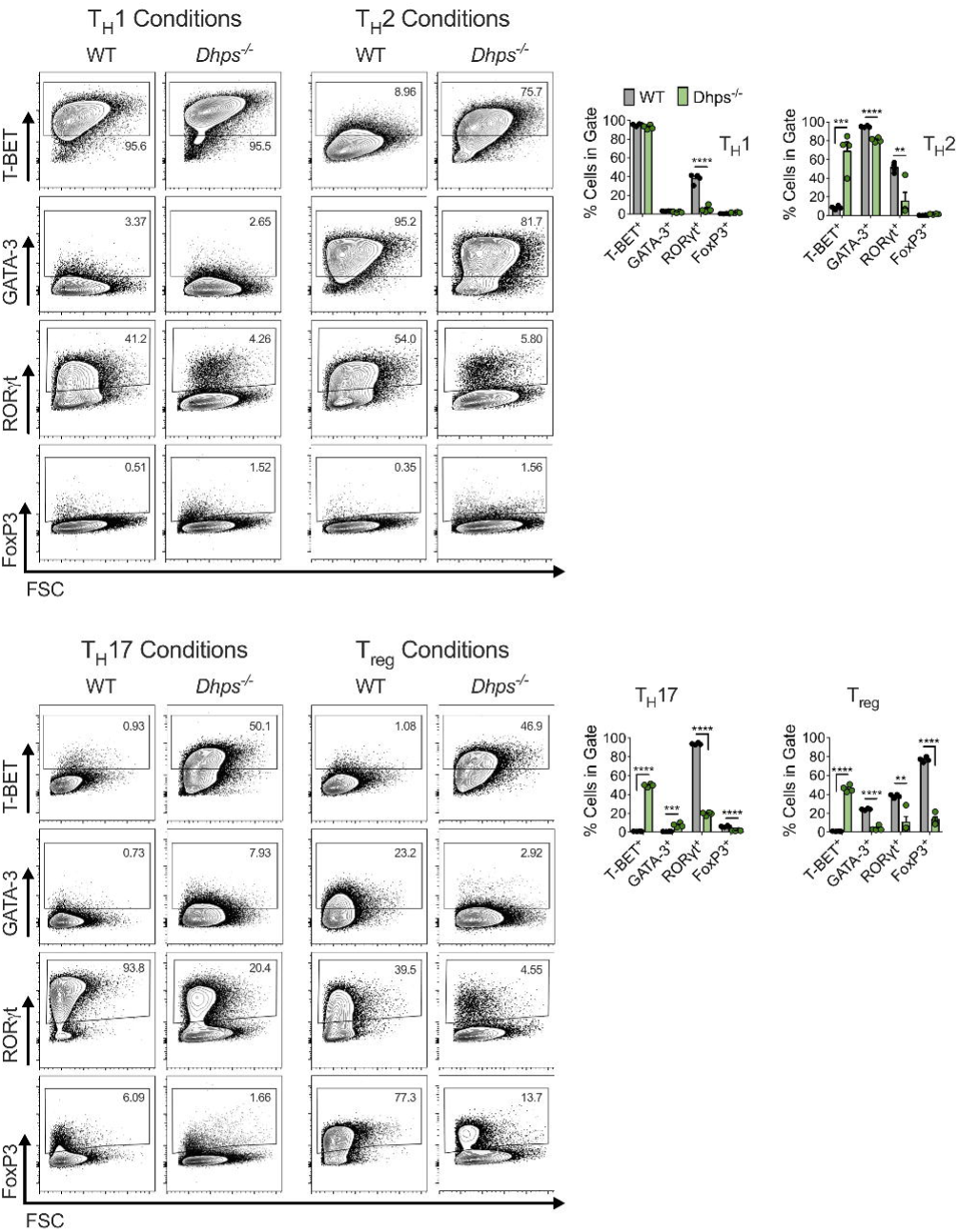
Transcription factor expression of *in vitro* activated *Dhps*^*-/-*^ CD4^+^ T cells (Related to Figure 4). WT and *Dhps*^-/-^ naïve CD4^+^ T cells were polarized for 4 days under indicated T_H_ cell conditions and the expression of labeled transcription factor assessed by flow cytometry. Representative contour plots are also shown. All data are mean ± SEM (p*<0.05, p**<0.005, p***<0.0005, p****<0.00005), representative of 3 experiments.

**Figure S8.**
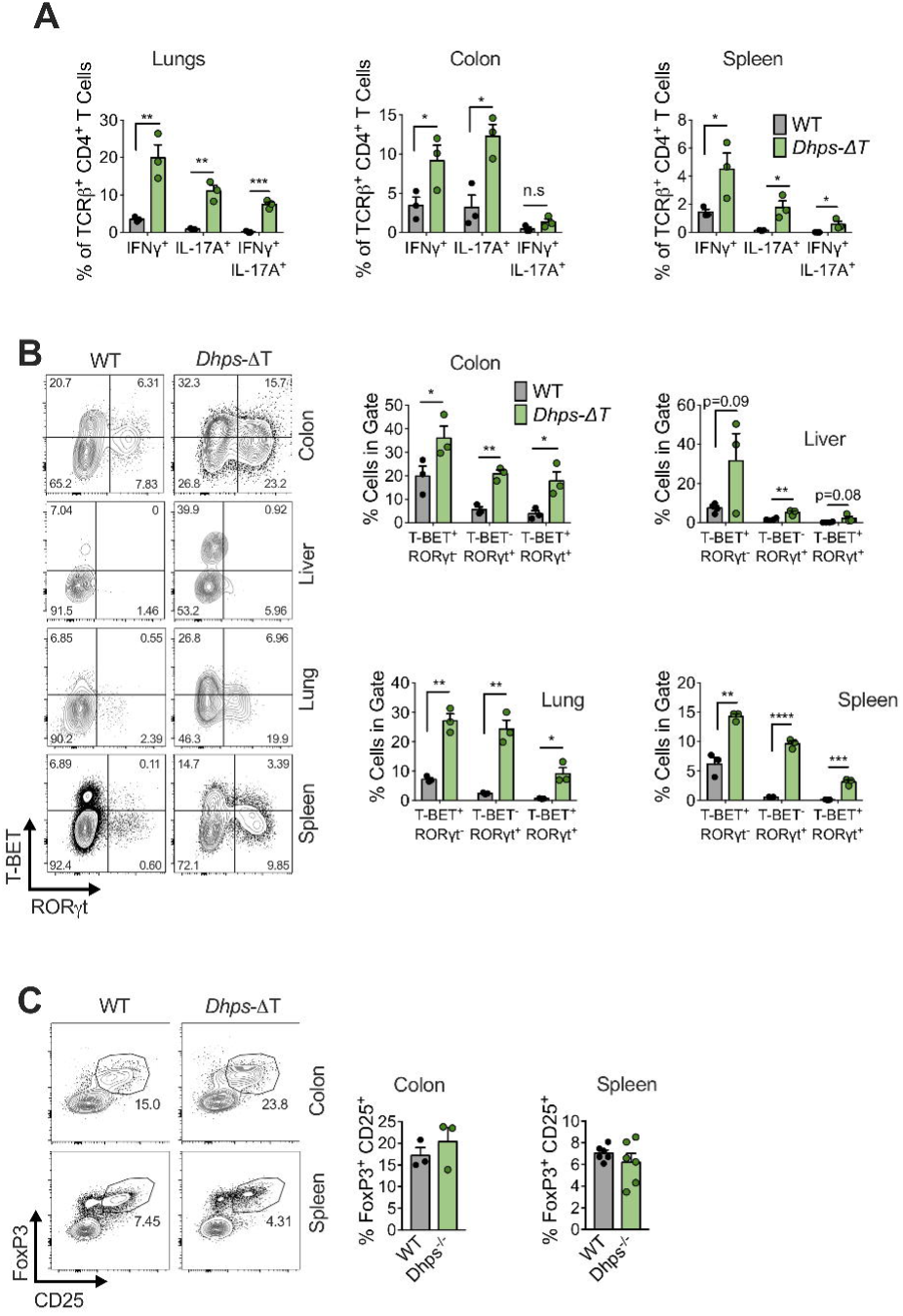
Cytokine and transcription factor expression of *Dhps*^*-/-*^ CD4^+^ T cells isolated *ex vivo* (Related to Figure 4). (**A**) CD4^+^ T cells were harvested from the designated organ of WT and *Dhps*-ΔT and cytokine expression was analyzed by flow cytometry following re-stimulated for 4 hours with PMA/ionomycin. (**B**) CD4^+^ T cells were harvested from the indicated organ of WT and *Dhps*-ΔT mice and the frequency of CD4^+^ T cells expressing labeled transcription factor was assessed by flow cytometry. Representative contour plots are shown. (**C**) Frequency of T_regs_ in CD4^+^ T cell compartment in colon and spleen of WT and *Dhps*-ΔT. All data are mean ± SEM (p*<0.05, p**<0.005, p***<0.0005, p****<0.00005). (A-C) Representative of 4 experiments.

**Figure S9.**
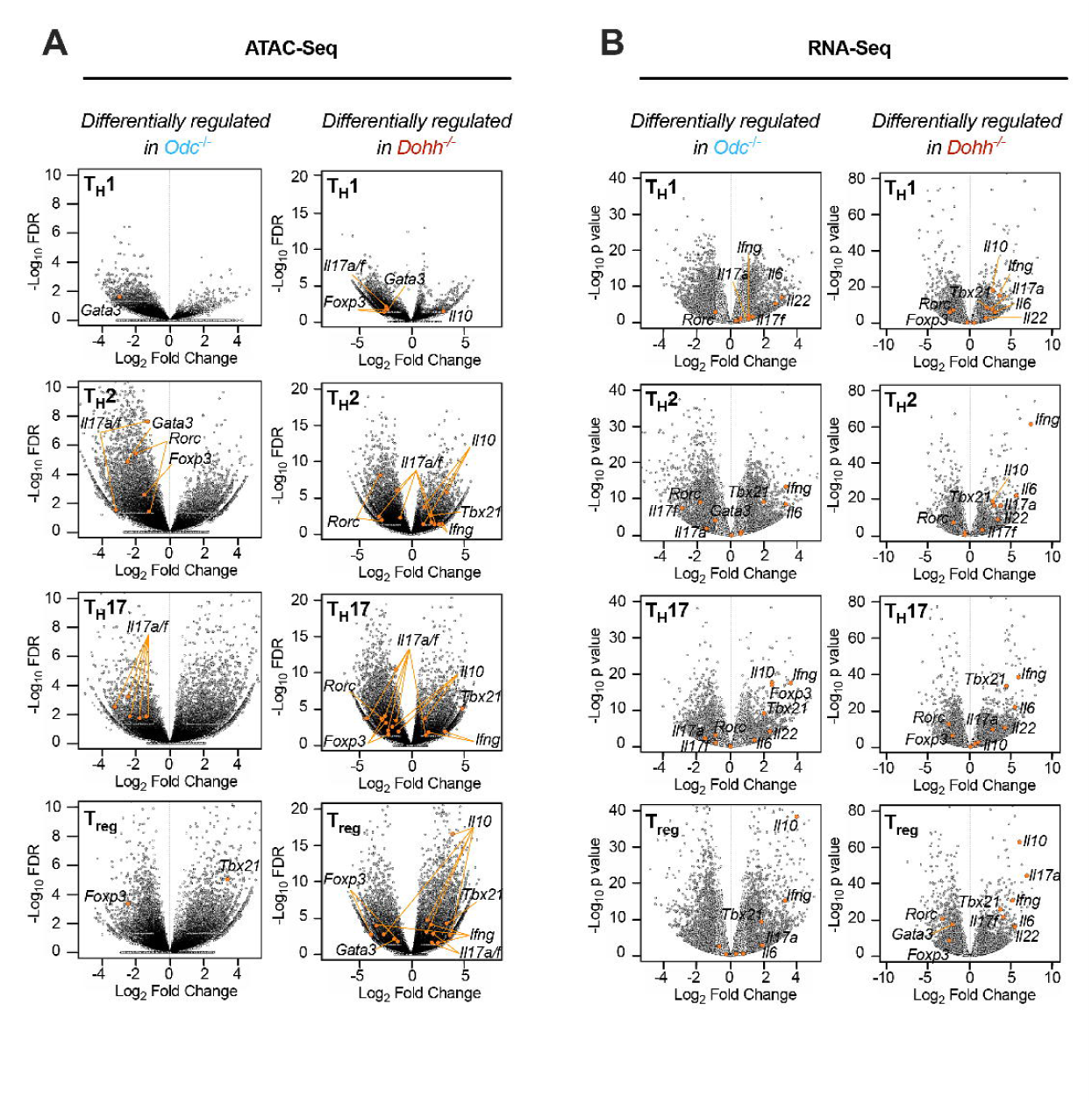
ATAC-Seq and RNA-Seq data from *in vitro* activated WT, *Odc*^*-/-*^, and *Dohh*^*-/-*^ CD4^+^ T_H_ cell subsets (Related to Figure 5). (**A**) ATAC-Seq of naïve CD4^+^ T cells isolated from the spleens and lymph nodes of WT, *Odc*-ΔT, and *Dohh*-ΔT mice that were activated with anti-CD3/CD28 for 4 days in indicated T_H_ cell polarizing conditions. Volcano plots depict all differentially regulated regions of open chromatin. **(B)** RNA was obtained and sequenced from naïve CD4^+^ T cells isolated from the spleens and lymph nodes of WT, *Odc*-ΔT, and *Dohh*-ΔT mice and polarized as in (A). Volcano plots depict differentially regulated genes in *Odc*^*-/-*^ and *Dohh*^*-/-*^ T_H_ cells relative to WT cells.

**Figure S10.**
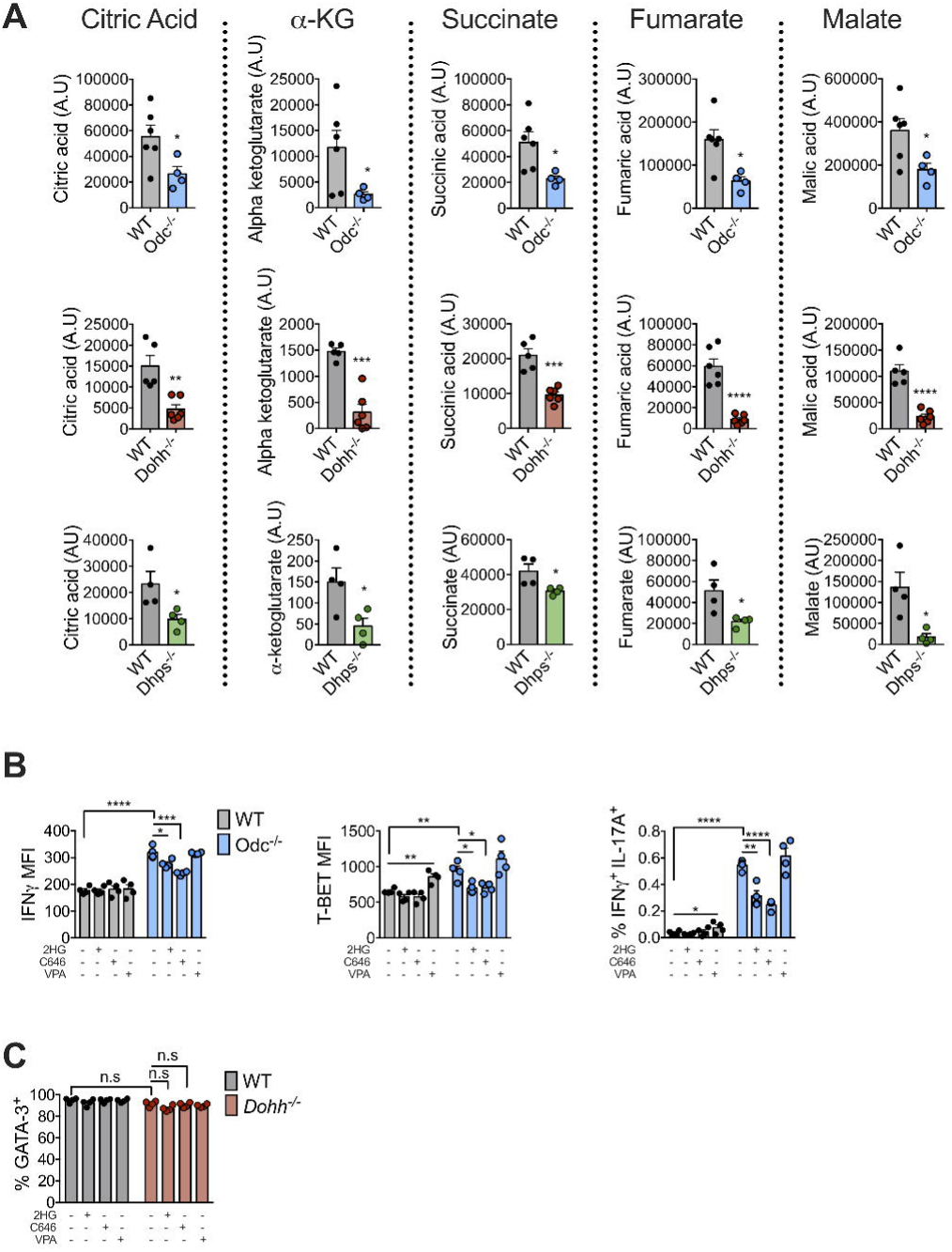
TCA cycle metabolites in activated *Odc*^*-/-*^, *Dohh*^*-/-*^, and *Dhps*^*-/-*^ CD4^+^ T cells and rescue of dysregulated transcription factor and cytokine expression (Related to Figure 5). (**A**) Naïve CD4^+^ T cells from *Odc*-ΔT, *Dohh*-ΔT, and *Dhps*-ΔT were activated with anti-CD3/CD28 for 48 hours and the quantity of each indicated metabolite was assessed by GC-MS. (**B**) Naïve CD4^+^ T cells from WT and *Odc*-ΔT mice were activated and polarized under T_H_2 conditions. After 24 hours cells were treated with either 25 mM D-2-HG, 20 μM C646 (HAT inhibitor), or 400 μM valproic acid (VPA, HDAC inhibitor) and the expression of indicated protein was assessed 48 hours later by flow cytometry. (**C**) Naïve CD4^+^ T cells from WT and *Dohh*-ΔT mice were activated and polarized under T_H_2 conditions. After 24 hours cells were treated with either 25 mM D-2-HG, 20 μM C646 (HAT inhibitor), or 400 μM valproic acid (VPA, HDAC inhibitor) and the expression of GATA3 was assessed 48 hours later by flow cytometry. All data are mean ± SEM (p*<0.05, p**<0.005, p***<0.0005, p****<0.00005). (A) Representative of 2 experiments, (B) Represents one experiment, (C) Representative of 4 experiments.

## METHODS

### Mice

Wildtype C57BL/6 and mice expressing Cre recombinase (CD4Cre) under the control of the CD4 promoter were purchased from Jackson Laboratories. *Dohh*^*flox/flox*^ and *Dhps*^*flox/flox*^ were a gift from Stefan Balabanov, Zurich. *Odc*^*flox/flox*^ mice were purchased from KOMP repository. All mice were bred and maintained under specific pathogen free conditions under protocols approved by the Animal Welfare Committee of the Max Planck Institute of Immunobiology and Epigenetics, Freiburg, Germany, and The St. Jude Institutional Animal Care and Use Committee, Memphis, USA, in accordance with the Guide for the Care and Use of Animals. Mice used for all experiments were littermates and matched for age and sex. Survival curves in Freiburg and Memphis were plotted using the time post-birth that mice were found with rectal prolopase, at which point mice would be sacrificed.

### Cell Culture

For in vitro culture, CD4^+^ T cells from the spleen and lymph nodes were isolated by negative selection using a Stem Cell kit according to the manufacturers instructions (Stem Cell, Cat: 19852). Cells were cultured in RPMI 1640 media supplemented with 10% FCS, 2 mM L-glutamine, 100 U/mL penicillin/streptomycin and 55 μM β-mercaptoethanol. All activations were done with 5 ug/mL anti-CD3 and 2 ug/mL anti-CD28 and 100 U/mL IL-2. T_H_ polarizations were performed as follows: T_H_1 – 4 μg/mL anti-IL-4, 10 ng/mL IL-12, 10 ng/mL IL-2. T_H_2 – 4 μg/mL anti-IFN-γ, 10 ng/mL IL-4, 10 ng/mL IL-2. T_H_17 – 10 μg/mL anti-IFN-γ, 10 μg/mL anti-IL-4, 5 ng/mL IL-6, 5 ng/mL TGF-β, 10 μg/mL IL-1β. T_reg_ – 4 μg/ mL anti-IFN-γ, 4 μg/mL anti-IL-4, 10 ng/mL TGF-β, 10 ng/mL IL-2. Cells were analyzed on day 4 of polarization. The following drug treatments were used where specified in the text: 20 μM C646, 25 mM D-2-hydroxyglutarate, 400 μM valproic acid, 250 μM putrescine hydrochloride (all Sigma).

### Western blot

For western blot analysis, cells were washed with ice cold PBS and lysed in 1 x Cell Signaling lysis buffer (20 mM Tris-HCl, [pH 7.5], 150 mM NaCl, 1 mM Na_2_EDTA, 1 mM EGTA, 1% Triton X-100, 2.5 mM sodium pyrophosphate, 1 mM β-glycerophosphate, 1 mM Na_3_VO_4_, 1 μg/mL leupeptin (Cell Signaling Technologies), supplemented with 1 mM PMSF. Samples were frozen and thawed 3 times followed by centrifugation at 20,000 x g for 10 min at 4°C. Cleared protein lysate was denatured with LDS loading buffer for 10 min at 70°C, and loaded on precast 4% to 12% bis-tris protein gels (Life Technologies). Proteins were transferred onto nitrocellulose membranes using the iBLOT 2 system (Life Technologies) following the manufacturer’s protocols. Membranes were blocked with 5% w/v milk and 0.1% Tween-20 in TBS and incubated with the appropriate antibodies in 5% w/v BSA in TBS with 0.1% Tween-20 overnight at 4°C. All primary antibody incubations were followed by incubation with secondary HRP-conjugated antibody (Pierce) in 5% milk and 0.1% Tween-20 in TBS and visualized using SuperSignal West Pico or femto Chemiluminescent Substrate (Pierce) on Biomax MR film (Kodak). Antibodies used: anti-ODC, anti-DHPS, anti-DOHH, anti-β-Actin (Abcam), anti-GAPDH (Cell Signaling), ant-EIF5A (BD Bioscience), anti-hypusine (Millipore).

### Polyamine Quantification by Mass Spectrometry

Metabolites were quantified by LC-MS using HILIC Chromatography on an Acquity UPLC BEH Amide column 1.7 µm, 2.1×100 mm on a 1290 Infinity II UHPLC system (Agilent Technologies) combined with targeted detection in a 6495 MS system (Agilent Technologies). Peak areas were normalized to ^13^C labelled internal standard (ISOtopic Solutions). Polyamines were extracted from 1 million CD4^+^ T cells using a mix of cold methanol, acetonitrile and water (50:30:20) containing 3% hydrochloric acid.

### Flow Cytometry

Flow cytometric staining was performed as previously described (Chang et al., 2015). To assess cytokine staining *in vitro* and *ex vivo*, cells were re-stimulated with 50 ng/mL PMA and 1 μg/mL ionomycin in the presence of brefeldin A (eBioscience) for 5 hours. Intracellular cytokine staining was performed using BD CytoFix/CytoPerm kit (BD Biosciences) and nuclear staining of transcription factors using the FoxP3 Permeabilisation kit (eBioscience). Cells were stained with Live/Dead viability dye (Thermo) prior to antibody staining. Cells were collected on LSR II and Fortessa flow cytometers (BD Biosciences) and analysed using FlowJo (TreeStar) software. The following antibodies were used: anti-CD4, anti-TCRβ, anti-CD45, anti-IL17A, anti-IL-17B, anti-IFN-γ, anti-T-bet, anti-GATA-3, anti-FoxP3 (all Biolegend) and anti-RORγt (BD Bioscience). For analysis of chromatin marks, cells were gated on Ki-67^+^ cells and diploid cells with ‘single’ DNA content based on FxCycle (dapi) staining in the live cell gate. Primary antibodies were stained for 90 minutes in permeabilization buffer at room temperature, followed by staining with the relevant secondary antibody for 30 minutes. The following antibodies were used: anti-H3k4Me^3^, anti-H3k9Ac, anti-H3k27Ac, anti-H3k9Me^3^ (all Cell Signaling), and anti-H3k36Me^3^ (Abcam).

### T cell transfer colitis

4×10^5^ naïve CD4^+^ T cells (CD45Rb^hi^ CD25^-^ CD44^lo^ CD62L^+^) from WT or *Odc*-ΔT mice were adoptively transferred into 12 week old *Rag1*^*-/-*^ recipient mice (Jackson) by intravenous injection. Disease score was calculated at the experimental endpoint using the following criteria:

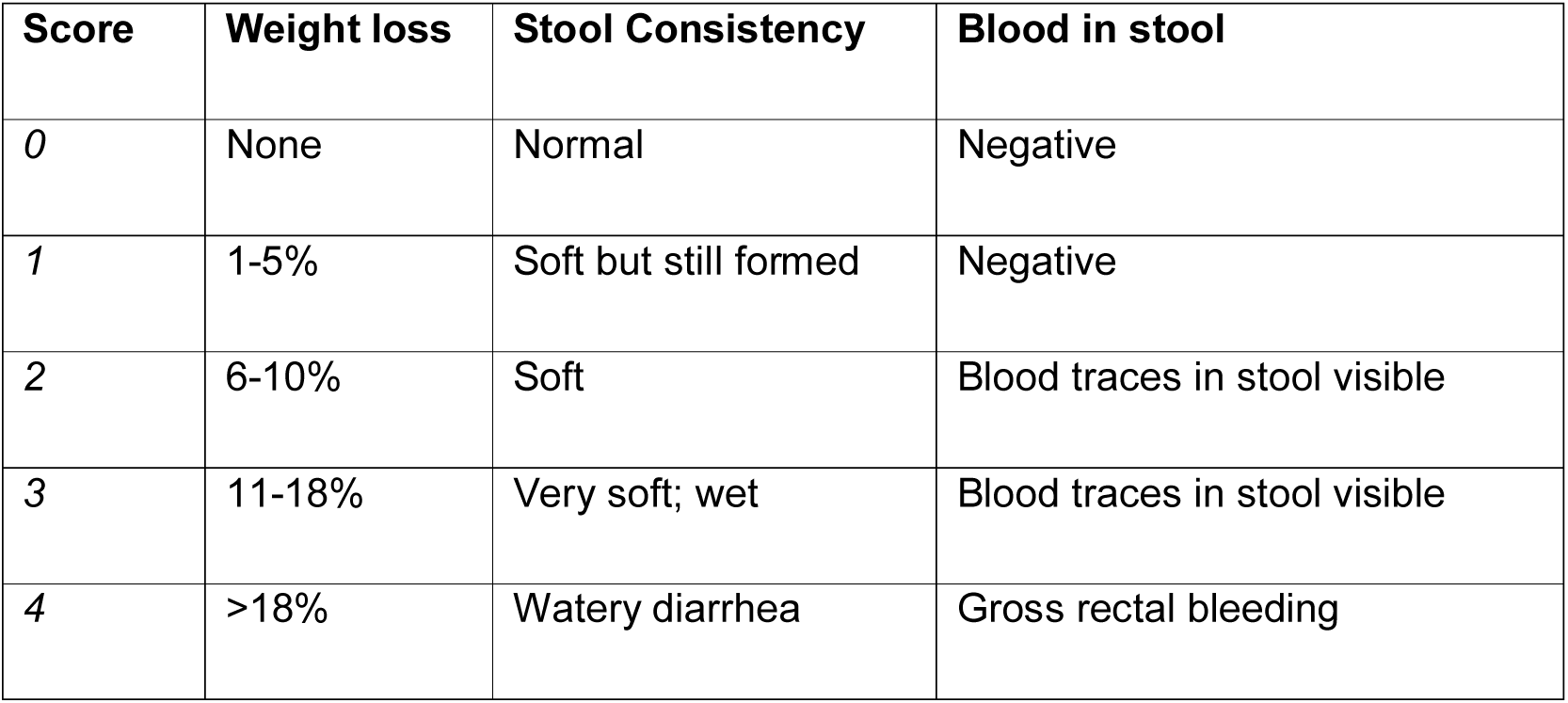

### *In vivo* treatment with anti-CD3 monoclonal antibody

Mice were injected intraperitoneally with CD3-specific antibody (clone 2C11, 50 µg/mouse) three times with 48 hours between each injection. Mice were sacrificed for analysis 4 hours after the third injection.

### Cytokine measurement in serum and supernatent

Serum and supernatent cytokine was measured by cytokine bead array using the LEGENDPlex Th cytokine panel according to the manufacturers instructions (Biolegend) on a BD Fortessa flow cytometer (BD Biosciences).

### Isolation of CD4^+^ T cells from Non-Lymphoid Tissues

For liver and lung, mice were perfused with PBS through the left ventricle. Organs were cut up into 1-3 mm^3^ pieces and digested in 1 ug/mL Collagenase A (Sigma) and 0.5 mg/mL Dnase I (Roche) in RPMI at 37°C for 30 minutes on a shaker. Digested organs were then finally mechanically disrupted through a cell strainer prior to flow cytometry staining. Cell suspensions from the colonic lamina propria were prepared as follows: Briefly, colons were isolated, cleaned, cut in small pieces and placed in in RPMI/5% FCS supplemented with 5mM EDTA. Tubes were then placed in 37°C in a shaking incubator to remove intestinal epithelial cells (IECs). This washing process was repeated twice, followed by one incubation with RPMI/5% FCS containing 15Mm Hepes. Digestion of the colon tissue was performed in RPMI/10% FCS containing 0.4mg/ml type VIII collagenase (Sigma Aldrich) and 40μg/ml DNase I (Roche) at 37°C for 60 minutes on a shaker. Supernatants were then filtered and a three-layered discontinuous Percoll gradient was used to obtain an enriched leukocyte fraction.

For the isolation of cells from the small intestine lamina propria – small intestines were isolated, cleaned and cut into 2cm pieces. Tissue was then incubated for 25 minutes in RPMI/3% FCS supplemented with 5mM EDTA and 0.15 mg/ml DTT at 37°C with shaking. After these small intestines were washed 3 times with RPMI containing 2mM EDTA. Tissue was then digested for 30 minutes in RPMI containing 0.1mg/ml Liberase TL (Roche) and 50ug/ml DNase I (Roche) at 37°C with shaking. After that a three-layered discontinuous Percoll gradient was used to enriched for the leukocyte fraction.

### Histology

Samples of the colon and caecum were collected and fixed in buffered 10% of 36% formalin solution for 24 hours and then stored in 70% ethanol prior to processing. Haematoxylin and eosin (H&E) staining was performed on 4–5 mm paraffin-embedded sections.

### RNA sequencing analysis

RNA was extracted using the RNeasy Kit (QIAGEN) according to manufacturer instructions and quantified using Qubit 2.0 (Thermo Fisher Scientific) following the manufacturer’s instructions. Libraries were prepared using the TruSeq stranded mRNA kit (Illumina) and sequenced in a HISeq 3000 (Illumina) by the Deep-sequencing Facility at the Max-Planck-Institute for Immunobiology and Epigenetics. Sequenced libraries were processed with deepTools (Ramírez et al., 2016), using STAR (Dobin et al., 2013), for trimming and mapping, and featureCounts (Liao et al., 2014) to quantify mapped reads. Raw mapped reads were processed in R (Lucent Technologies) with DESeq2 (Love et al., 2014) to generate normalized read counts to visualize as heatmaps using Morpheus (Broad Institute) and determine differentially expressed genes with greater than 2 fold change and lower than 0.05 adjusted p value. Gene ontology analysis was performed used the free online platform DAVID (Da Wei Huang et al., 2009) and Ingenuity® pathway Analysis (QIAGEN). Supervised clustering of gene expression was performed with pheatmap (version2012) using Ward’s minimum variance method (Murtagh and Legendre, 2014).

### ATAC sequencing analysis

Libraries were prepared using the Nextera DNA library Prep Kit (Illumina) adapting a published protocol (Buenrostro et al., 2015). Briefly, 5×10^4^ T cells treated as described were washed in PBS and then lysed in 10 mM Tris-HCl, pH 7.4,10 mM NaCl, 3 mM MgCl_2_ and 0.1% Igepal CA-630 (all Sigma). Nuclei were then spun down and then resuspend in 25 μl TD (2x reaction buffer), 2.5 μL TDE1 (Nextera Tn5 Transposase) and 22.5 μL nuclease-free water, incubated for 30 min at 37°C. DNA was purified with the Qiagen MinElute PCR Purification Kit (Thermo Fisher Scientific). PCR amplification was performed with the NEBNext High-Fidelity 2x PCR Master Mix (New England Labs) using custom Nextera PCR Primers containing barcodes. Adaptors were removed with AMPure XP beads according to manufacturer’s protocol. Libraries were quantified with the Qubit and submitted for sequencing with a HISeq 3000 (Illumina) by the staff at the Deep-sequencing Facility at the Max-Planck-Institute for Immunobiology and Epigenetics. Sequenced samples were trimmed with Trimmomatic ^9^ and mapped using Bowtie2 (Ben Langmead and Salzberg, 2012). Open chromatin was detected with MACS2 (Zhang et al., 2008), while differences between treatments was determined using DiffBind (Ross-Innes et al., 2012) with at lest 2 fold change in accessibility and a false discovery rate lower than 0.05. For visualization only, replicate mapped files were merged with SAM tools ^13^ and coverage files were generated with deepTools and visualized alongside coverage files on IGV (Robinson et al., 2011). Bed files were analyzed with Bedtools ^15^.

### Statistical Analysis

p-values were determined using unpaired Student’s t-test. Differences were considered statistically significant when p<0.05 (*<p0.05, **p<0.01, ***p<0.001). Data are shown as mean ± s.e.m. Statistics were calculated using GraphPad Prism 6 software.

